# Divergent Visuomotor Strategies in Teleosts: Neural Circuit Mechanisms in Zebrafish and *Danionella cerebrum*

**DOI:** 10.1101/2024.11.22.624938

**Authors:** Kaitlyn E. Fouke, Zichen He, Matthew D. Loring, Eva A. Naumann

## Abstract

Many animals respond to sensory cues with species-specific coordinated movements to successfully navigate their environment. However, the neural mechanisms that support diverse sensorimotor transformations across species with distinct navigational strategies remain largely unexplored. By comparing related teleost species, zebrafish (*Danio rerio, ZF*) and *Danionella cerebrum* (*DC*), we investigated behavioral patterns and neural architectures during the visually guided optomotor response (OMR). Closed-loop behavioral tracking during visual stimulation revealed that larval *ZF* employ burst-and-glide locomotion, while larval *DC* display continuous, smooth swimming punctuated with sharp directional turns. Although *DC* achieve higher average speeds, they lack the direction-dependent velocity modulation observed in *ZF*. Whole-brain two-photon calcium imaging and tail tracking in head-fixed fish reveals that both species exhibit direction-selective motion encoding in homologous regions, including the retinorecipient pretectum, with *DC* exhibiting fewer binocular, direction-selective neurons overall. Kinematic analysis of head-fixed behavior reveals that *DC* sustain significantly longer directed swim events across all stimuli than *ZF*, highlighting the divergent visuomotor strategies, with *ZF* reducing tail movement duration in response to oblique, turn-inducing stimuli. Lateralized motor-associated neural activity in the medial and anterior hindbrain of both species suggests a shared circuit motif, with distinct neural circuits that independently control movement vigor and direction. These findings highlight the diversity in visuomotor strategies among teleost species, underscored by shared sensorimotor neural circuit motifs, and establish a robust framework for unraveling the neural mechanisms driving continuous and discrete visually guided locomotion, paving the way for deeper insights into vertebrate sensorimotor functions.

**Research Highlights:**

- *Larval DC* exhibit faster swimming than *ZF*, matching the direction of visual motion.
- *DC* execute OMR in smooth, curved swimming patterns, interspersed with sharp directional turns.
- *ZF* and *DC* share similar visuomotor neural architecture, recruiting pretectal and hindbrain regions.
- *ZF* and *DC* demonstrate lateralized encoding of turns, particularly in medial hindbrain neurons.

**In Brief:** Larval *Danionella cerebrum* respond to global visual motion cues in smooth, low-angle swimming patterns, interspersed with sharp directional turns, swimming consistently faster than zebrafish. Fouke et al. use behavioral tracking of freely moving and head fixed fish to reveal an evolutionarily conserved visuomotor neural architecture transforming visual motion cues into species-specific locomotor behaviors.

**Graphical Abstract.**
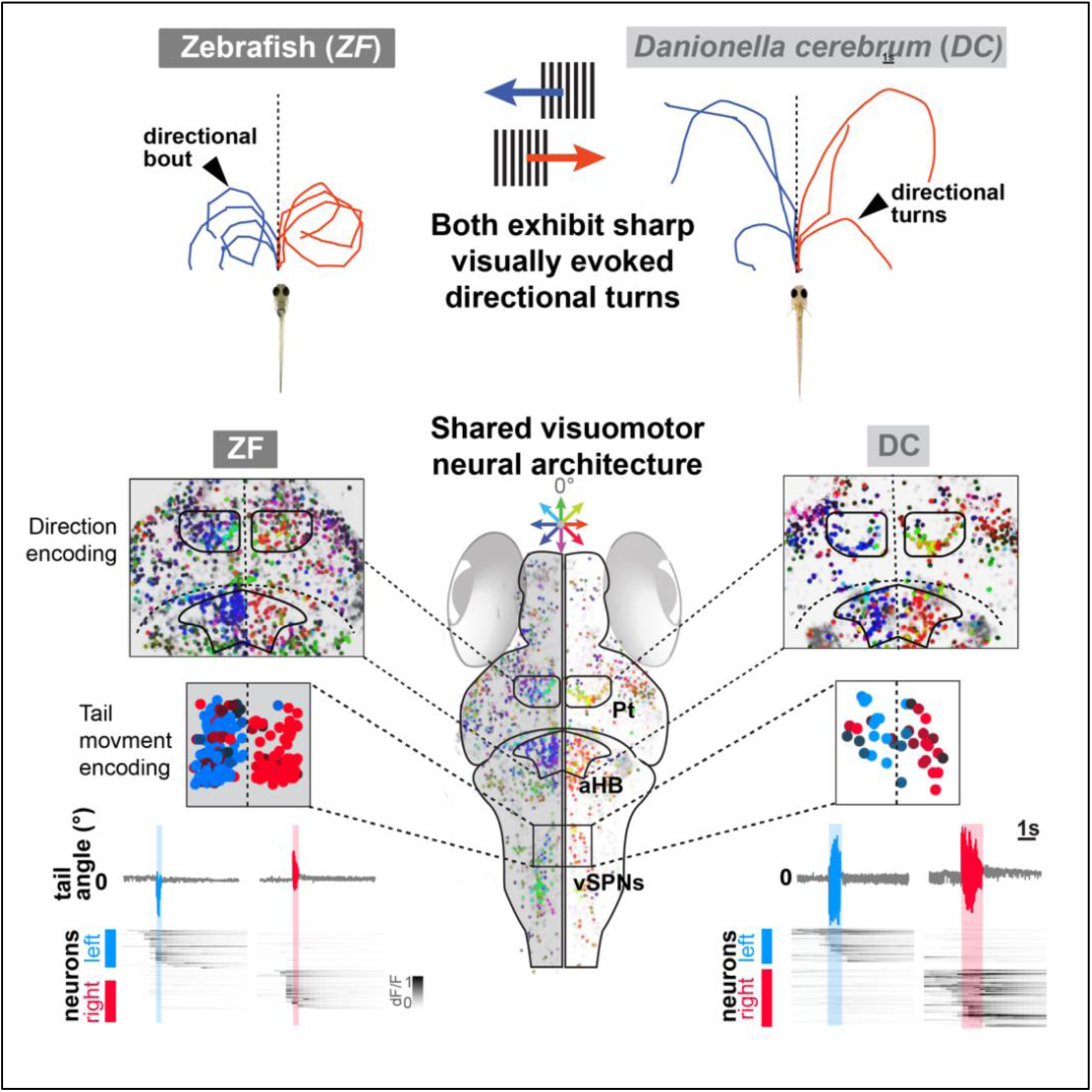

## Introduction

To successfully navigate their environment, animals frequently respond to sensory input with discrete, directed movements—with a clear start and end^1^. For instance, catching a moving object requires the brain to convert constant visual motion cues into different motor outputs of specific speed and duration^2^. One common visuomotor transformation is the optomotor response (OMR), which stabilizes the body and retinal image by following the moving visual scene induced by displacements in water or wind currents, present across many species, from birds^3^, fruit flies^4^ to fish^5^, and even humans^6^. Across vertebrates, the underlying neural systems processing the visual information in the retina^7^ and generating the locomotion output are highly conserved^8^. In aquatic environments, the OMR is critical in helping animals stabilize their body position in water currents by compensating for displacement through optic flow processing^9^. To investigate the neural circuit activity underlying OMR behaviors, recent studies have exploited the optical and genetic accessibility of the translucent larval zebrafish (*Danio rerio*)^10–12^. However, the prevalence of OMR across teleost species with different ecological niches, behavioral strategies, and neural computations remains unclear.

Recently, the related teleost species, *Danionella cerebrum* (*DC*), commonly known as the micro glassfish, has attracted interest as a neuroscience vertebrate model owing to its small brain volume^13^, life-long transparency^13–16^, and unique ecological niche^17,18^. These miniature cyprinids exhibit various social behaviors, including acoustic communication with the capacity for directional hearing^19^, socially reinforced learning^20^, social behavior development^21^, and social place preference^22^. While larval zebrafish (*ZF*) naturally inhabit clear, shallow water, *DC* are found in deeper water levels with dimmer visual environments^17,18,23,24^. During spontaneous locomotion, larval *ZF* swim in discrete bouts, characterized by alternating short phases of undulating tail movements and passive glides^24,25^. In contrast, larval *DC* spontaneously move by swimming continuously using long, consistent tail beats, exhibiting long ballistic, straight swimming phases with a persistent heading direction^24^. Remarkably, these species exhibit distinct visually triggered startle responses, with *DC* and *ZF* responding more to dark-to-light and light-to-dark transitions, respectively^26^. The experimental advantages shared by *DC* and *ZF*^13,24^ provide a unique opportunity to investigate species-specific behavioral adaptations and underlying neural circuit computations.

During OMR, 5-day-old larval *ZF* robustly follow global optic flow by executing routine turns and engaging the retinorecipient pretectum (*Pt*), which contains spatially intermixed, directionally tuned neurons that show sustained responses to visual motion stimuli^10,27,28^. In *ZF*, these *Pt* neurons anatomically connect to downstream motor command regions, including identifiable descending spinal projection neurons of the midbrain nuclei of the medial longitudinal fasciculus (*nMLF*)^29^. These *nMLF* neurons show intermittent, bout-like firing patterns and preferentially respond to tail-to-head (forward) motion, which drives increased bout frequency in freely swimming *ZF*^30,31^. To execute directional turns, *ZF* initiate bouts with a strong tail bias (i.e., a higher curvature) to either side^32^. A different set of identifiable ventromedial spinal projection neurons (*vSPNs*) in the medial hindbrain activate ipsilateral motor neurons to initiate these biased turns^32^. These *vSPNs* show functional lateralization, with *vSPN* neurons on the left preferring left visual motion and vice versa^32^. Unilateral ablation of these *vSPNs* in *ZF* suggested that these descending neurons are necessary to adjust the direction of the first tail-beating amplitude^32^. While *DC* exhibit a homologous set of these spinal projection neurons^24^, their function has not been characterized in the context of visually evoked behaviors. More broadly, we lack insight into how *DC* respond to visual motion or how their neurons process visual cues to guide locomotion.

In this study, we compare *DC* and *ZF* to uncover the neural mechanisms underlying their natural variation of visuomotor coordination. Using closed-loop behavioral tracking in freely swimming fish and tail tracking during volumetric two-photon calcium imaging, we characterize species-specific visuomotor responses and map the brain-scale neural circuitry that encodes visual motion and locomotion, revealing shared visuomotor architectures with distinct adaptations. This comparative framework highlights both conserved and divergent aspects in natural variations of teleost visuomotor pathways, offering insights into how species-specific adaptations shape neural processing and behavior.

## Results

### Larval DC exhibit higher average swimming speeds during OMR

To investigate OMR behaviors in larval *DC* compared to *ZF*^10–12^, we conducted high-speed, closed-loop behavioral tracking while projecting visual motion stimuli from below. In an automated trial structure, individual fish were exposed to moving gratings (10 mm/s) in whole-field cardinal, intercardinal, and eye-specific right and left directions (**Figure 1A, S1A**). Despite the similarity in size and shared morphological features between larval *DC* and *ZF* (**Figure 1B**), their locomotion patterns differ considerably^24^. While *ZF* execute sequences of discrete locomotion bouts consisting of clear burst and glide phases, *DC* swim continuously^24^. Consistent with previous studies^12,30^, *ZF* respond vigorously to forward, tail-to-head (0°) moving gratings, increasing forward bout frequency to match the optic flow direction. These visually evoked bouts in *ZF* consist of sharp velocity increases (peak velocity = 44.7 mm/s, median velocity = 1.8 mm/s) followed by pauses in movement, resulting in a periodic pattern of velocity peaks and interbout glides. We demonstrate that *DC* also perform OMR behaviors, but they do so by following the optic flow direction with continuous steady swimming, maintaining a relatively constant velocity (median velocity = 13.7 mm/s, **Figure 1C**). Unlike the discrete burst-and-glide movements of *ZF*, which allow clean extraction and analysis of individual bouts^10,12,24,30,33,34^, *DC*’s continuous swimming style presented challenges for identifying distinct behavioral events (**Figure S2**). Therefore, to compare OMR behavior across *ZF* and *DC,* we calculated instantaneous behavioral kinematics, e.g., velocity (distance swum per second), angular velocity (orientation change per second), and angular acceleration (orientation change per second squared). While *ZF* increased their average velocity during forward motion presentation (8.8 mm/s) compared to stationary gratings (5.0 mm/s, Student T-test, p < 0.01), *DC* did not significantly upregulate their velocity during forward motion (forward = 9.7 mm/s, static = 10.6 mm/s, Student T-test, p = 0.89, **Figure 1D**). Similarly, when comparing OMR performance to eight directions of whole field motion, *ZF* strongly modulate their average velocity depending on the motion direction, e.g., with backward motion resulting in decreasing velocity (4.3 mm/s). In contrast, *DC* maintained a consistently higher average velocity across stimuli than *ZF* (*DC* = 9.8 mm/s, *ZF* = 6.4 mm/s ANOVA, p > 0.05, **Figure 1D**).

**Figure 1.**
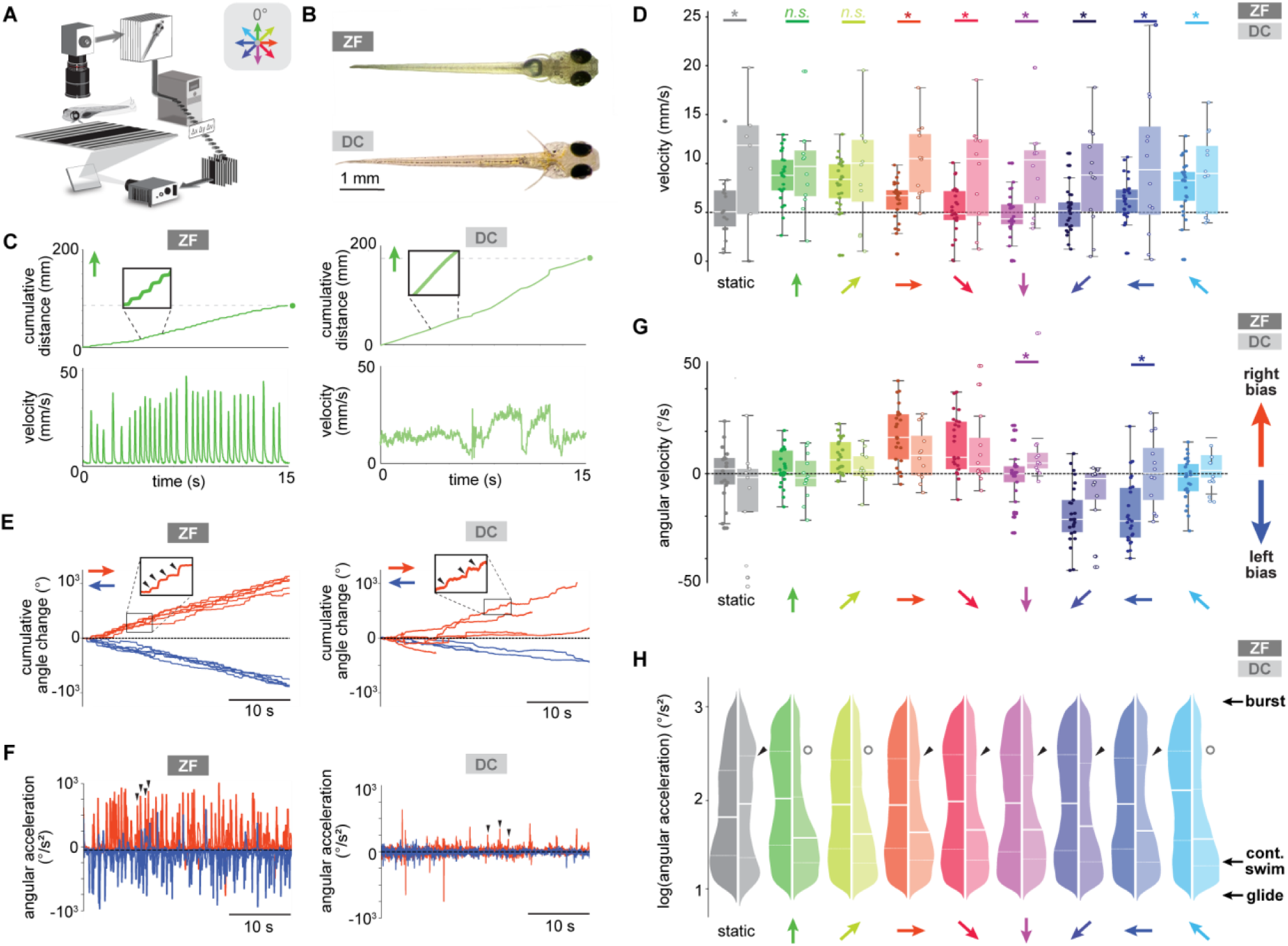
Freely swimming *DC* perform OMR with smooth, continuous swims, punctuated with sharp turns. **A** Closed-loop behavioral assay to track freely swimming fish gratings in response to cardinal and intercardinal directions. Heading direction (Δν) and position (Δx, Δy) are extracted to lock stimuli to the body axis. **B** 7-day-old zebrafish (*ZF*) and 8-day-old *Danionella* cerebrum (*DC*). **C** Cumulative distance and velocity of representative trials for *ZF* and *DC* to forward motion (green). *ZF* move in discrete movements, while *DC* swim continuously, resulting in DC traveling farther over 15 sec (green circles). **D** Median velocity for *ZF* (N = 26) and *DC* (N = 12) during visual stimulation. While *ZF* exhibit direction-dependent velocity modulation, *DC* consistently swim at a higher median velocity than *ZF* across stimuli, without modulating velocity. (p> 0.05, two-way ANOVA). **E** Cumulative heading direction change to rightward (red) and leftward motion (blue) of representative trials for *ZF* and *DC*. *Zoom in*: sharp orientation changes (arrowheads) were observed in both species. **F** Instantaneous angular acceleration for the same trials as in **E**. For leftward and rightward motion, *ZF* generates many sharp turns, *DC* move in ballistic, straight swims with occasional directed, high-angle turns (arrowheads). **G** Median angular velocity for data in **D**. Both species follow the direction of visual motion, with positive and negative angular velocity indicating rightward and leftward turning, respectively. Asterisks indicate a significant difference between *ZF* and *DC* angular velocity (p> 0.05, two-way ANOVA). **H** Instantaneous angular accelerations for *ZF* and *DC* for same experiments in **D**, **G**. Logarithmic distributions highlight differences in movement modes. High angular acceleration values indicate sharp, burst-like turns lower values occur during continuous, smooth turns (> 10 °/s^2^) or slower, gliding. Center white lines indicate median, dashed lines quartiles. All *ZF* and *DC* angular accelerations are significantly different (p> 0.05, Mann-Whitney U test). *ZF* swim mostly in bouts (higher acceleration, black arrows).

### Optic flow induces angular matching and turns in DC

When presented with stimuli orthogonal to the body axis, such as left or rightward moving gratings, *ZF* responded with characteristic consecutive routine turns in the same direction as the stimulus^10,12^. For instance, *ZF* executed leftward bouts in response to leftward moving gratings (**Figure 1E**). As the stimulus remained locked to the fish’s body axis, *ZF* displayed routine turns during oblique motion stimulation (>90°), increasing their angular velocity (°/s) and exhibiting rapid, instantaneous angular acceleration (°/s^2^, **Figure S1C**). In comparison, larval *DC* executed OMR in smooth, curved trajectories interspersed with sharp directional movements (**Figure 1E**, black arrowheads). During oblique motion stimulation, *DC* accelerated quickly during these punctuated directional turns while maintaining continuous tail undulation with fluctuations in angular acceleration to complete broad turns (**Figure 1F**). While *DC* exhibited predominantly low-angle trajectories to align with the stimulus direction (**Figure 1G**), they repeatedly executed sharp angle turns in response to directional stimuli, resulting in brief, high angular acceleration during oblique motion trials (**Figure S1B**, **Video 1**).

During all motion cues, *ZF* maintained higher angular acceleration than *DC*, as *ZF* move in discrete bouts (*ZF* = 87.8 °/s^2^, *DC* = 41.6 °/s^2^, **Figure 1H**), with periods of lower angular acceleration events indicating a decrease in acceleration during the glide phase. Exposure to oblique motion revealed that *DC* can generate sharp turns like *ZF*, displaying increased instantaneous angular acceleration events during these stimuli (**Figure 1H**). Notably, even during the static control condition, when stationary gratings were locked to the fish body, *DC* showed higher angular acceleration, suggesting that the high contrast visual cues alone can induce sharp turns in *DC*. To probe the contribution of each eye to the OMR, we also presented *ZF* and *DC* with monocular left and rightward moving stimuli (**Figure S3A**). As *ZF*, *DC* moved in the stimulus direction but generally responded less to monocularly presented stimuli, indicating that *DC* require more visual motion to drive OMR behaviors (**Figure S3B**). Together, these comparative kinematic analyses demonstrate that, as *ZF*, larval *DC* can overall perform the required neural processing to perform OMR behaviors. Nonetheless, *DC* present reduced turning, angular acceleration, and binocular integration, and lacks the stimulus direction-dependent modulation of their speed observed in ZF.

### DC and ZF share a conserved visuomotor neural architecture

To examine visually evoked neural activity in *DC* and *ZF,* we performed whole-brain calcium imaging with volumetric, two-photon microscopy (**Methods**, **Video 2**). Both species, expressing *GCaMP7f* across their brain, were head-fixed in agarose while visual motion stimuli were projected from below (**Video 3**). To link neural activity with behavior, we displayed the same stimuli, correlating fluorescence changes with different directions of motion (**Figure 2A**). Using unbiased source extraction^35^, we identified direction-selective units of the size of the average zebrafish neuron (diameter = ∼7 um^36^), from here on referred to as neurons, that consistently responded to repetitions of the same motion stimuli (**Figure 2B**). At the population level, *ZF* and *DC* exhibited neurons across the brain that responded to all directions of motion (**Figure 2C**). The spatial distributions of these direction-selective neurons reveal that motion information is distributed in the lateralized retinorecipient pretectum and the anterior hindbrain for *ZF* and putative homologous regions in *DC* (**Figure 2D**, **S4**). The hierarchical clustering of motion responses further indicated that both species share major sets of neurons with specific response profiles (**Figure 2D, S5**). Despite the similarity in neuroanatomical distribution and direction selectivity across neurons in both species, *DC* contained fewer motion-responsive neurons compared with *ZF* (average DSI: *ZF* = 0.62, *DC* = 0.66, p > 0.05, Student’s t-test; neural unit count: *ZF* = 12249, *DC* = 4841; p < 0.05, Student’s t-test, **Figure 2E, F**). The lower number of motion-responsive neurons, along with a greatly reduced average number of forward-tuned neurons (forward-tuned neurons: *ZF* = 1494, *DC* = 562, p < 0.05, Student’s t-test), cannot be explained by lower brain volume in *DC* (average brain volume: *ZF* = 5.2 * 10^7^ um^3^, *DC* = 4.7 * 10^7^ um^3^, p > 0.05, Student’s t-test). The reduced prevalence of these neurons may reflect adaptations to distinct environmental niches with varied levels of visibility and possibly explain *DC*’s reduced velocity modulation during forward OMR in freely swimming fish.

**Figure 2.**
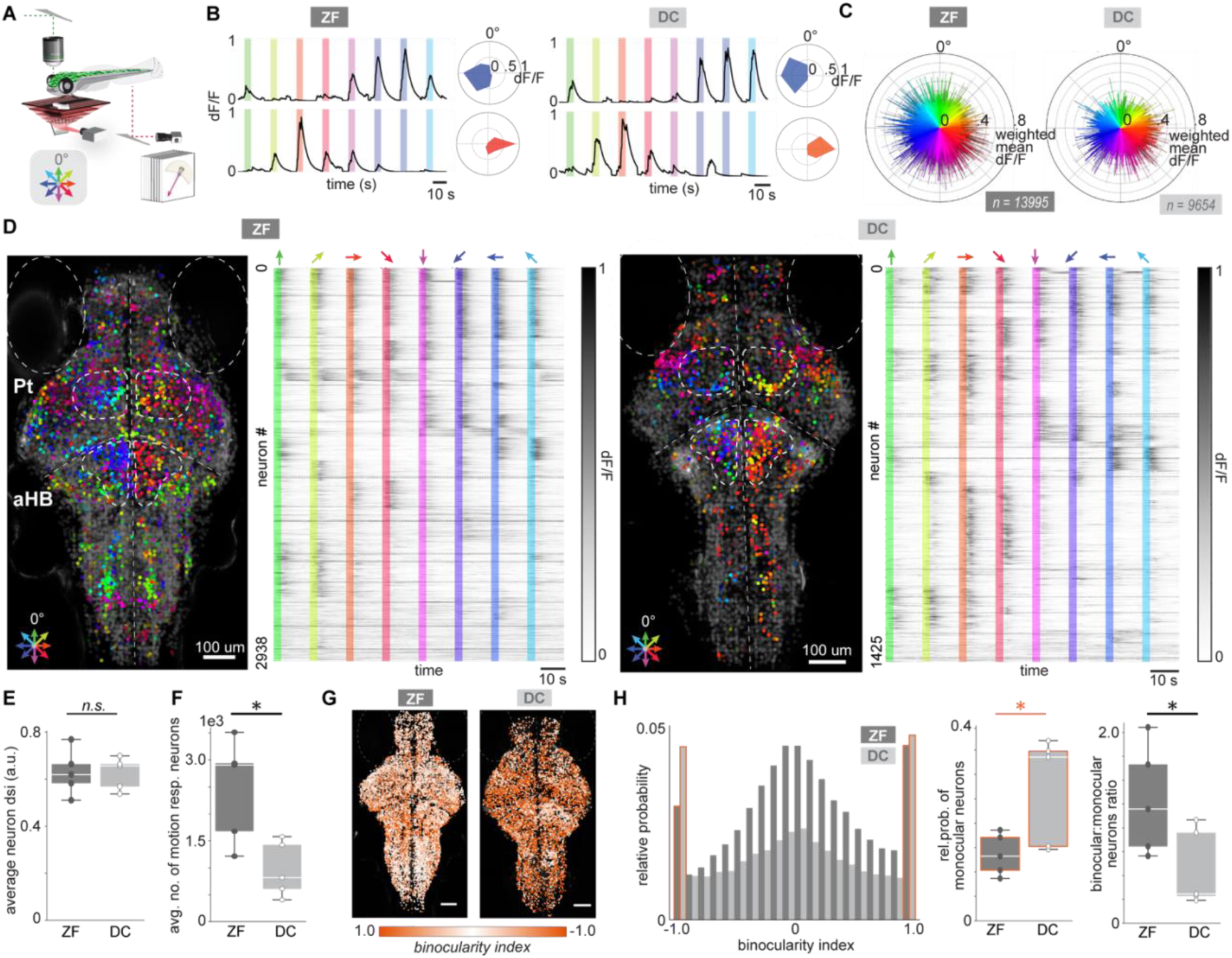
Larval *DC* and *ZF* share visual motion processing neural architecture. **A** Imaging setup. Volumetric two-photon calcium imaging of head-fixed, tail-freed larval fish during visual stimulation (color wheel). **B** Average dF/F activity of representative rightward and leftward direction-selective *ZF* and *DC* neurons, with associated rose plots, color-coded for direction selectivity. **C** Representative vector plots of all motion-responsive neurons in *ZF* (N = 13995/17445 total detected neurons) and *DC* (N = 9654/12134 total detected neurons). Vector angle represents direction selectivity index (DSI), length indicates weighted mean, and neurons with mean response <0.2 dF/F are shaded grey. **D** Whole-brain distribution of visually responsive neurons for data in **C**. Each dot represents a neural source, its hue indicates direction selectivity (≥ 0.2 mean dF/F), otherwise grey, plotted on a greyscale *GCaMP* image. Homologous pretectum (Pt) and anterior hindbrain (aHB) regions in *ZF* and *DC* contain lateralized direction-selective neurons*. Right,* hierarchical clustering of trial averaged dF/F of direction-selective neurons with mean response >0.2 dF/F across both species (N = *ZF*: 2938, *DC*: 1425, ***Methods***). **E** Average DSI is similar in *ZF* and *DC* (p > 0.05, student t-test; fish N = *ZF*: 5, *DC*: 5 fish). **F** *ZF* exhibit more motion-responsive neurons than *DC* (p < 0.05, student t-test; N = *ZF*: 5, *DC*: 5 fish). **G** Binocularity index (BI) maps for representative *ZF* and *DC*. **H** Histogram of BI distribution of all motion-responsive neurons in *ZF* and *DC* (N = *ZF*: 55324, *DC*: 36049 neurons). *DC* show a higher proportion of purely monocular neurons (p > 0.05, student t-test; N = *ZF*: 5, *DC*: 5 fish) than *ZF*. BI ratios reveal a higher fraction of binocular neurons in *ZF* compared to *DC* (p < 0.05, student t-test; N = *ZF*: 5, *DC*: 5 fish).

### DC relies on more monocular-tuned neurons

To investigate binocular integration at the neural level^10^, *DC* and *ZF* were presented with left and rightward monocular and binocular motion stimuli (**Methods**). By calculating a binocularity index (BI) for each neuron to determine activation by the left or right eye, we generated volumetric brain maps showing the distribution of monocular and binocular neurons (**Figure 2G**, **S4**). Consistent with previous ocular classification in *ZF*^10,27,37,38^, both species contain many monocular neurons, which respond exclusively to motion in one eye, and binocular neurons, which respond to motion in both eyes (**Figure 2H**). However, *DC* exhibited a significantly higher proportion of monocular neurons compared to *ZF* (relative probability: *ZF* = 0.13, *DC* = 0.34, p < 0.05, Student’s t-test), and fewer binocular neurons (BI ratio: *ZF* = 1.19, *DC* = 0.31, p = 0.05, Student’s t-test). This increased reliance on monocular processing in *DC* likely limits its ability to integrate visual information across both eyes, conceivably contributing to its reduced OMR compared to *ZF*, where more neurons facilitate binocular integration to drive locomotion.

### Head-fixed DC performs OMR with longer swim events than ZF

To link visually evoked neural activity with behavior in head-fixed *DC* and *ZF*, we analyzed tail movements during calcium imaging (**Figure 3A**). Using highspeed (200 Hz) infrared video to track 13 points along the tail (**Video 3**), we captured individual swim events by summating the angles of each tail segment (**Figure 3B**, **S6A**). Visual motion stimuli reliably triggered locomotion swim events in both species (**Figure S7**), but while *ZF* exhibited short, discrete swim patterns aligned with the direction of motion, *DC* showed prolonged swim events, adjusting the direction via multiple larger tail directional movements within a single swim event (arrowheads, **Figure 3B**, **S6B**). Consistent with velocity control differences in freely swimming fish, unlike *DC*, *ZF* modulated swim event duration based on stimulus direction, significantly longer swims for forward stimuli and shorter for oblique stimuli (**Figure 3C**, **S7A**, **B**). Both species matched stimulus direction in this head-fixed configuration, but *ZF* exhibited significantly greater directional turning than *DC* (**Figure 3D**). While both species showed similar tail beat frequencies and tail angle distributions (**Figure S7D**, **E**), *DC* displayed significantly longer swim events across all stimuli than *ZF* (**Figure 3C, E, F**, **S7A**). These results indicate species-specific differences in locomotion persistence and shared direction adjustments are preserved even in head-fixed conditions.

**Figure 3.**
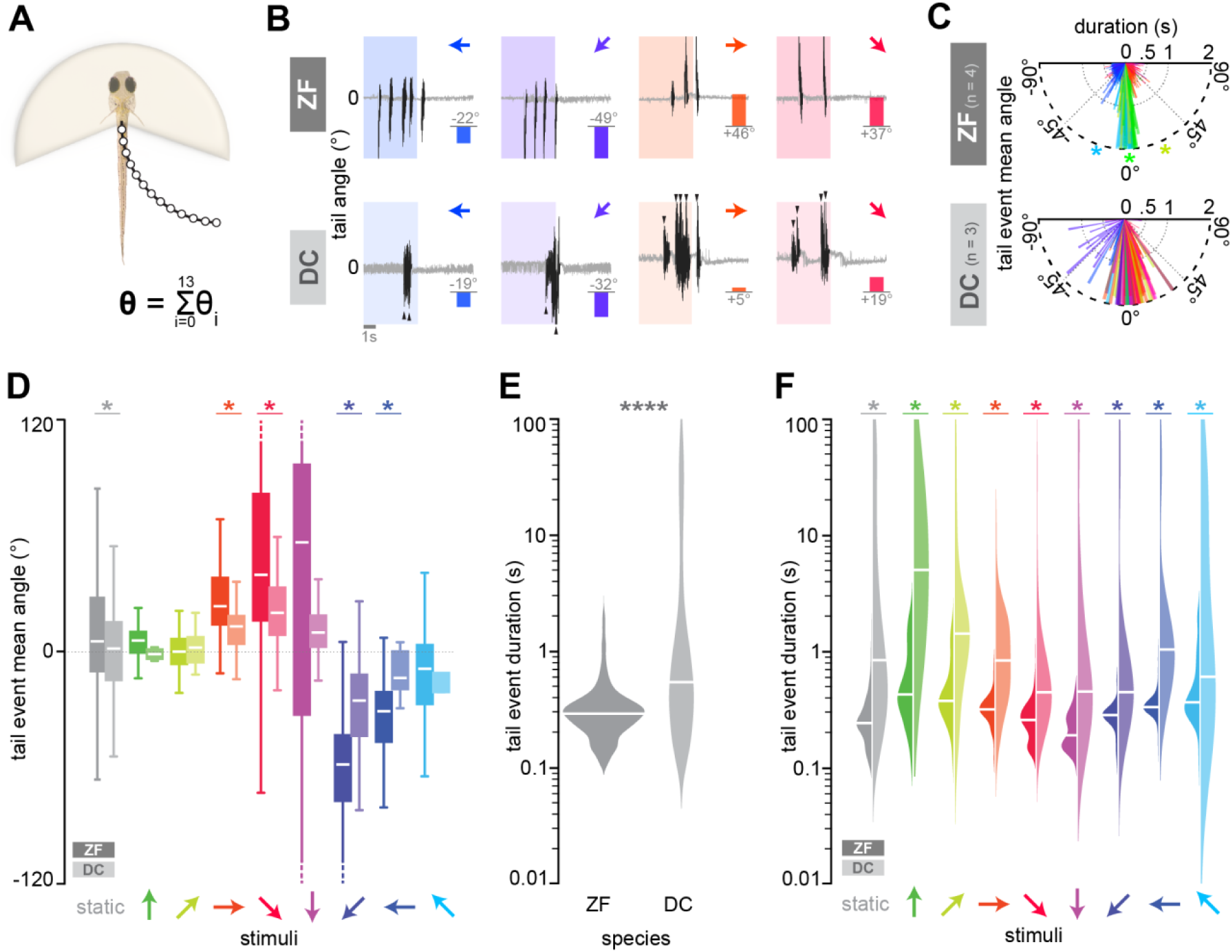
Head-fixed larval *DC* exhibit longer visually evoked swim events than *ZF.* **A** Schematic of tail tracking in head-fixed fish during two-photon calcium imaging. The tail angle (Θ) is the sum of all angles between each segment. Positive and negative Θ indicate right and left tail movements. **B** Representative *ZF* and *DC* tail events during visual motion (colored shaded areas, arrows indicate direction). Black traces represent tail angles during detected events, otherwise grey. Arrowheads highlight large tail flicks in *DC*. Bar graphs show the mean tail angle magnitude, and the direction averaged across each event during motion presentation. **C** Distribution of all visually evoked tail events in head-embedded *ZF* (N = 4 fish, 1887 events) and *DC* (N = 3 fish, 331 events). Each bar represents a tail event, with its direction indicating tail angle and color matching the visual stimulus direction (color wheel). Bar length encodes tail event duration, capped at 2s. In *ZF*, tail events during forward stimuli last significantly longer than those during turn-inducing stimuli (* p < 0.05, two-way ANOVA, Tukey post hoc). *DC* tail events are less lateralized, with no significant shortening for oblique stimuli. **D** Mean tail angle across all tail events for *ZF* and *DC*, white lines indicate the medians. Both *DC* and *ZF* turn in the optic flow direction, with *ZF* exhibiting higher angle turning than *DC* during turn-inducing stimuli (* p < 0.05, two-way ANOVA, Tukey post hoc)**. E** Violin plots with medians (white lines) of all tail event durations for *ZF* (N = 4 fish, 3568 events) and *DC* (N = 3 fish, 855 events) plotted on a log scale. *DC* perform significantly longer swims than *ZF* (**** p < 0.0001, two sample t-test). **F** Distribution of all tail event durations, white lines indicate medians. *DC* show significantly longer tail events than *ZF* (* p < 0.05, two-way ANOVA, Tukey post hoc)

### Motor-associated activity supporting species-specific locomotion

To elucidate the neural dynamics driving visually evoked behavior in *DC* and *ZF*, we analyzed neurons with activity correlated with tail movements. We hypothesized that calcium dynamics in relevant neurons would predict motor output, and, conversely, that motor output would predict neuronal calcium activity (**Figure 4A**, **Methods**). Using this bi-directional decoding approach, we observed neurons with elevated calcium activity during swim events, revealing bout-locked activation patterns for shorter *ZF* swims and repeated activity throughout the duration of *DC*’s longer swims (**Figure 4B**, **S8A**, **B**). Across all fish, motor-associated neurons were consistently found in the midbrain *nMLF* region^12,24,30^, anterior hindbrain, known for influencing turning ^33,39^, and throughout the hindbrain (**Figure 4C**, **S8A**). Among these, the top 20% of motor-associated neurons were enriched in these homologous brain regions, with several temporally close bouts in *ZF* and sustained swims in *DC* driving elevated calcium activity in these neurons (**Fig S8C**, **D**), indicative of increased neural activity during vigorous or frequent events. Additionally, longer swims in *ZF* and stronger tail bends in *DC* correlated with more robust population responses (**Figure S8E**, **F**). General Linear Model (GLM) analysis revealed that in *DC,* maximal tail angle predominantly drives motor-associated neural dynamics (**Figure S8F**). In contrast, in *ZF,* swim duration is a key factor, highlighting adaptive neural mechanisms for their distinct locomotion strategies even in head-fixed fish. Across all motor-associated neurons, *ZF* primarily showed brief, bout-associated neural activity, while *DC* consistently displayed characteristic ongoing neural activity, tiling across prolonged swim events (**Figure 4D**).

**Figure 4.**
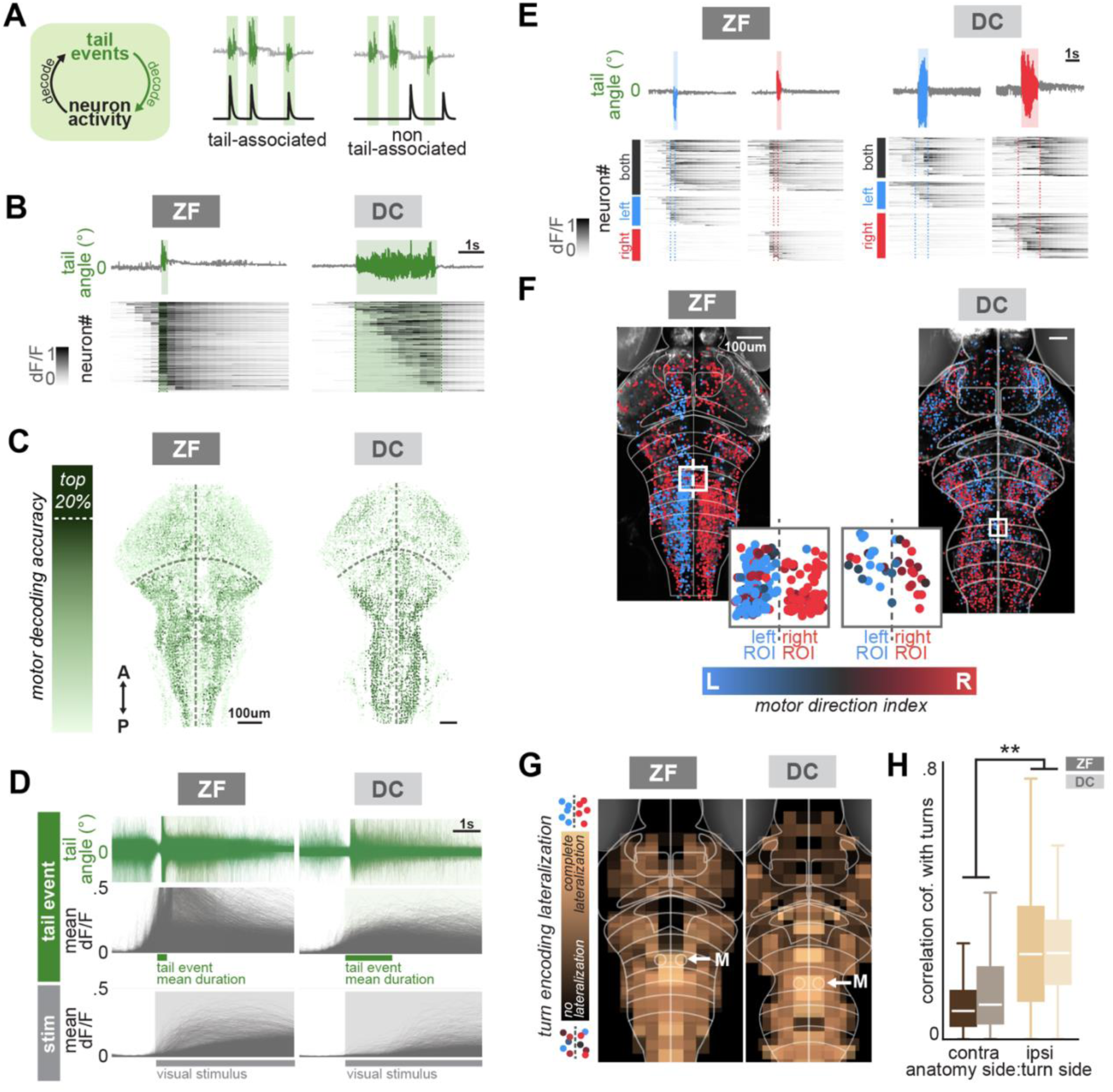
Medial hindbrain neurons encode turning behaviors in both species. **A** Schematic of the bidirectional decoding approach to identify motor-associated neurons based on the temporal alignment of calcium signal increases with tail events, quantifying each neuron’s motor decoding accuracy. *Right*, idealized illustrations of motor-associated and non-motor-associated neurons. **B** Average dF/F responses of highly motor-associated neurons during representative tail events for *ZF* and *DC*. Green shading denotes the tail movement duration. **C** Anatomical distribution of motor decoding accuracy of all neurons in representative *DC* and *ZF* brain volumes, highlighting enrichment of motor-associated neurons in the hindbrain and *nMLF* regions. Darker colors indicate higher motor decoding accuracy. **D** The top 20% of motor-associated neurons are not responsive to visual stimuli (N = *ZF*: 4 fish, 14063 neurons; *DC:* 2 fish, 5790 neurons). *Top*, aligned tail events (green). *Middle*, average calcium activity aligned to the start of all tail movement events. *Bottom*, aligned to visual stimulus motion onset. **E** Top 20% motor-associated neurons during representative left and right turns in *ZF* and *DC*, showing neurons selectively associated with left, right, or both turning directions. **F** Distribution of the turning direction encoding by top 20% motor-associated neurons in *ZF* and *DC*. Inset: neurons near rhombomere 4 (white box) exhibit strongly lateralized activity, with each neuron color-coded by its motor direction index, i.e., neurons on the anatomical left respond more during left turns and vice versa. **G** Spatial map illustrating the degree of turning encoding lateralization across *ZF* and *DC*. Rhombomeres surrounding Mauthner cells (M) display the strongest lateralization (N = *ZF*: 4 fish, *DC*: 2 fish)**. H** Within the medial hindbrain (boxed ROI in **F**), motor-associated neuron activities correlate more with the motor regressor representing ipsilateral turning. Boxplots represent Pearson’s correlation coefficients for motor-associated neurons in the ROIs across fish (N = *ZF*: 4 fish, 707 neurons; *DC*: 2 fish, 112 neurons, (** p < 0.01, two-way ANOVA).

### Lateralized encoding of turning behaviors in medial hindbrain neurons in both species

To identify motor-associated neurons involved in directional turns, we applied a similar bi-directional decoding algorithm to distinguish neurons selectively responsive to left or right tail movements (**Figure 4E**, **S9A**, **B**). Each neuron was assigned a motor direction index, calculated as the average mean tail angle across concurrent swim events, weighted by the magnitude of neural activity (**Methods**). While motor direction indices varied across brain regions in individual fish (**Figure 4F**), neurons in the medial hindbrain area, specifically located within rhombomeres 4 to 6, displayed strong, lateralized encoding of turning direction in both species (**Figure 4G**). This region, identified by locating Mauthner neurons in both species (**Figure S9C**), contained turn-encoding neurons that anatomically co-localize with the identifiable *vSPNs*^32^. Computing a lateralization index further demonstrated that these neurons activate preferentially during left or right turns to the corresponding side (**Figure 4H**). Together with distinct turning patterns in freely swimming *DC* and *ZF* (**Figure 1H**, **S1**), these results indicate that both species share homologous neural circuit mechanisms that control directed movements and neural activity that supports species-specific differences in swim vigor and duration in visuomotor transformations.

## Discussion

Behavioral diversity, crucial for species-specific survival, arises from complex neural circuits transforming sensory inputs into motor outputs. Yet, the evolutionary changes in neural function driving this diversity remain largely unexplored^40,41^. In this comparative study, we leverage two teleost species, *Danionella cerebrum* (*DC*) and zebrafish (*ZF*), to characterize the neural circuits that process visual motion to drive divergent locomotion patterns. First, we show that both species execute visuomotor behaviors to align themselves with the optic flow direction. While *ZF* display strong stimulus direction-dependent velocity modulation, *DC* swims continuously in straight, ballistic trajectories, achieving faster average speeds than *ZF* (**Figure 1**). Despite these differences in locomotion strategies, we demonstrate that *DC* adjust direction via interspersed sharp turns, similar to *ZF*’s well-characterized routine turns during OMR^42^. Second, whole-brain imaging indicates both species rely on spatially intermixed populations of directionally tuned neurons in overall conserved, homologous brain regions^10,27^, including pretectum and anterior hindbrain (**Figure 2**). Third, we demonstrate that the species-specific visuomotor behavioral phenomena can be investigated during whole-brain calcium imaging (**Figure 3**). Specifically, we identified motor-associated neurons in similar brain areas that control overall locomotion and support the shortened and prolonged locomotion patterns in *ZF* and *DC*, respectively. Further, we show that lateralized encoding of turns in the medial hindbrain region drives directional tail movements in both species. The prevalence of these turn-encoding neurons, colocalized with the reticular spinal projection neurons, underscores a conserved mechanism for directional control during visuomotor tasks (**Figure 4**). This work establishes the experimental and analytical framework to compare visually evoked behaviors and underlying neural circuitry across species with natural circuit variations. By identifying conserved and divergent elements in the visuomotor pathways of *DC* and *ZF*, our findings provide a foundation for understanding how species-specific adaptations influence neural processing and behavior.

### Continuous swimming versus burst and glide OMR

Our study reveals that larval *DC* respond to visual motion by swimming continuously, consistent with observations from previous findings^24^, contrasting larval *ZF*, which typically exhibit burst-like routine turns^25,42^. This continuous swimming may reflect adaptations to *DC*‘s low-light, high-turbidity, and low-oxygen habitats^17^, where visual input is less reliable and constant movement aids oxygen intake^24^, suggesting ecological and foraging strategies as key factors shaping their distinct locomotion patterns. In contrast, *ZF* may rely on discrete bouts to improve visual processing by reducing self-generated visual input, which might benefit navigation in clear, high-contrast environments^34,43–45^. While our behavioral characterization demonstrates that larval *DC* are fully capable of executing OMR behaviors, approximately only half of all tested *DC* triggered the complete set of our OMR trials, possibly because of their tendency towards thigmotaxis^26^. Potentially, other experimental features in our assays, such as too bright, high contrast, and fast-moving visual stimuli, may cause behavioral inhibition, or *DC*’s continuous swimming may be too energy-consuming^46–48^. Since *DC* has been characterized as a highly social species ^19–22^, the stress of social isolation may cause increased thigmotaxis or negative phototactic drive. Studying *DC* OMR with varying stimulus conditions or in a school of *DC* fish could provide insights into the modulation of visuomotor behavior and the role of social reinforcement.

Next, confirming *DC*’s OMR capacity, we uncovered that visual stimuli drive sharp directional turns characterized by angular tail movements and large heading direction changes, riding on top of the characteristic continuous swimming (**Figure 1H**). Exposed to directional visual stimuli, *DC*’s sharp turns aid in rapidly aligning the fish’s body axis, stabilizing retinal image and body position in perceived optic flow. Burst-like turning surpasses broad, smooth turning as the more energetically advantageous behavior strategy^49^. Mechanistically, sharp turning could present an advantageous strategy to respond quickly to changes in visual scenery or moving objects. This observation points to the explanation that *DC* and other continuous swimming teleost species, such as medaka^50,51^, are capable of rapid directional adjustments in response to external sensory input, such as looming^52^, OMR^53^ or prey stimuli^50^.

### Neural circuitry variations in DC and ZF

Despite these behavioral differences, the anatomical distribution of visually responsive neurons is conserved in 6-9 days post fertilization (dpf) *DC* and *ZF*. In both species, visually responsive neurons are located in the retinorecipient pretectal regions and the anterior hindbrain, areas well-characterized in the processing of whole-field visual motion^10,27,33,39^. However, *DC* exhibit overall fewer motion-responsive neurons, relatively more monocular-tuned neurons, possibly contributing to *DC*’s reduced propensity for OMR behaviors. By allocating fewer neurons to visual processing and functional shift towards monocular motion processing could present a general adaptation to low light conditions^57^. This characteristic aligns with the hypothesis that continuous forward motion in *DC* relies less on dynamic binocular adjustments, potentially reflecting an energy-efficient adaptation for habitats with limited visual cues^54^. Similar shifts in monocular versus binocular processing have been observed in species adapted to dimly lit or turbid environments, underscoring how environmental demands can recalibrate sensorimotor processing pathways^44,50,55^. It is plausible that the cost of transportation (COT) of continuous swimming mode is more energy-consuming, and therefore, whenever we observed *DC* swimming, modulation was not necessary. Similarly, it could suggest a lack of forward-tuned visual input to the neurons of the *nMLF*^24^. Nonetheless, the otherwise strong directional tuning of visually responsive neurons in *DC* suggests no reduction in qualitative visual processing but likely a different behavioral interpretation of these visual signals.

By simultaneous behavioral tracking and neural activity recordings across both species of brains, we could identify species-specific directional tail movements and large populations of motor-associated neurons. In *ZF*, turns are driven by the spinal projection neurons distributed between rhombomere 4-6^12,32^. Consistent with our freely swimming analysis, *DC* contains neurons in these homologous areas, strongly correlated with directional tail events (**Figure 4F**). These results suggest that these descending vSPNs drive visually evoked turns in both species. This underscores the shared circuit motif for abrupt directional adjustments riding on top of general locomotor drive that generates bouts in *ZF*, and prolonged swims in *DC*. Therefore, we propose that *ZF* and *DC* regulate swim vigor and duration via symmetric medial locomotion maintenance neurons (MLMNs) that drive continuous swimming^24^, yet engage homologous visually responsive descending functionally lateralized *vSPNs* to adjust direction. This combinatorial circuit independently controls swim vigor and direction, concurrently controlling tail undulations and modulating turning bias. As previously suggested^24,56^, the species-specific spinal cord central pattern generators potentially contribute to the differences in *ZF* and *DC* swim duration, employing specific stop neurons to terminate tail undulation phases. These overall conserved sensorimotor neural architectures suggest that, although *DC* and *ZF* have evolved different behavioral strategies to cope with their respective environments, they retain fundamental homologies in the neural circuits that govern visuomotor integration.

### Visuomotor processing variations in fish

Our characterization of OMR behavior and neural circuitry in *DC* establishes the foundation for exploring the natural circuit variations, which has been successfully used in comparing various physiological features in seeing surface and blind cave dwelling *Astyanax mexicanus*^57^. Other continuous swimming teleost fish, like the medaka, have also been shown to detect and capture prey with one eye^50^, further supporting this shift to monocular processing in *DC*. As we observed relatively few forward-tuned neurons, the paucity of these neurons may explain why *DC* does not modulate its velocity when presented with forward motion. Other fish, including sharks, with poorly lit habitats also show a lower optokinetic gain^58^, a related phenomenon to whole body OMR. Although other fish species, such as medaka, goldfish, guppy, three-spined stickleback, mbuna, glowlight tetra, and bronze Corydoras, perform OMR behaviors^59^, *DC* and *ZF* offer experimental tractability to comprehensively map and ultimately understand evolutionary-driven behavioral diversity and the underlying neural circuitry.

In conclusion, our findings highlight the value of *Danionella cerebrum* as a comparative model for studying how homologous neural architectures can give rise to divergent visuomotor strategies. Future investigations could expand to causally test the neural mechanistic underpinnings of these behaviors, including optogenetic manipulations, which could provide a deeper understanding of how evolution shapes species differences^60,61^ in visuomotor transformation. Our findings not only deepen our understanding of species-specific adaptations in visuomotor control but also highlight shared, conserved neural circuit motifs, suggesting that diverse locomotor patterns can emerge from similar neural architectures adapted to each species’ needs. Together, we demonstrate the capacity of OMR behaviors in another teleost species driven by a conserved functional neural architecture.

## Methods

### Zebrafish and Danionella cerebrum husbandry

Live-fish experiments were approved by Duke University School of Medicine’s standing committee of the animal care and use program (IACUC). *ZF* (*Danio rerio)* and *DC* (*Danionella cerebrum*) colonies were raised in the Duke University School of Medicine Zebrafish Core Facilities in Durham, NC. All adult fish were grown at a 14:10 hour light/dark cycle at a water temperature of ∼27.5°C, pH 7.2, and 500 μS conductivity, with fewer than 20 adult fish per tank. Adult *ZF* were fed Gemma Micro 150 and live, enriched *Artemia* once per day. *ZF* embryos were collected from adult breeder zebrafish and groups of ∼30 fish raised in Petri dishes with E3 egg medium. Once the fish reached four days post fertilization (dpf), they were fed live *Paramecia* once a day. Similarly, adult *DC* were fed Gemma Micro 150 once per day and live, enriched *Artemia* twice per day. Because *DC* spontaneously spawn in crevices^13^, 2-3 2-inch PVC pipes were added to each tank. <25 adult fish per tank. The larval *DC* were grown at a density of <15 larvae per Petri dish in E3 medium and incubated at 28.5°C. They were fed live *Paramecia* once per day between 4-7 dpf and fed both *Paramecia* and Z Plus after 7 dpf. Behavior and imaging experiments were performed on larval fish 5-9 dpf at ages when sex cannot be determined. Before imaging, we screened both species for bright fluorescence, indicating high *GCaMP* expression to achieve the best calcium signals. In general, the *DC* adults and larvae were significantly more delicate than the *ZF* adults and larvae, requiring careful handling. For free swimming behavioral experiments, we used larval Ekkwill (EK) *ZF*, wildtype *DC*; for imaging *Tg(elavl3:GCaMP7f) ZF* and *Tg(neurod:GCaMP7f) DC*. GCaMP7f *ZF* were a generous gift from Dr. Misha Ahrens. Wildtype *DC* were graciously provided by Dr. Adam Douglass, and *Tg(neurod:GCaMP7f) DC* by Drs. Lovett-Baron, Ahrens, and colleagues at Janelia Research Campus.

### Freely swimming optomotor response behavior

To measure visually evoked behaviors in freely swimming larval *ZF* or *DC*, we used six closed-loop behavioral rigs in parallel to run fully automated experimental OMR routines adapted from previous work^10^. To orchestrate trial-based behavioral tracking and presentation of visual stimuli (**Figure 1A**), we integrated our custom *pandastim* visual stimulus software with the Python-based open-source *Stytra* package^62^. Briefly, we projected visual stimuli from below a behavioral arena while tracking their x,y position and orientation at >150 Hz using a high-speed tracking infrared-sensitive CMOS camera (FLIR, GS3-U3-41C6NIR-C: 4.1 MP, 90 FPS, CMOSIS CMV4000-3E12, NIR) with a 35mm 1” lens (Edmund Optics, 35mm Focal Length Lens, 1” Sensor Format, #63-247). We illuminated the fish from below with custom infrared (850 nm) illuminator panels (CMVision, CM-IR130-850NM) paired with an 830 nm long-pass filter (Edmund Optics, SCHOTT RG-830). Visual stimuli were displayed onto a diffusing screen below the behavioral arena via reflection from a cold mirror (both Edmund optics) with a P300 Pico Projector (AAXA technologies). As previously published^10^, we implemented a trial structure starting with the presentation of concentrically moving rings to orient and guide individual fish toward the center of the Petri Dish. The detection of the fish in the center triggered the start of an OMR trial. Each trial began with a 3-second control phase in which stationary gratings would be locked to the fish’s body position before the gratings started moving at 0.12 mm/sec. Whole field moving gratings were presented at angles of 0°, 45°, 90°, 135°, 180°, 225°, 270°, or 315° relative to the fish’s orientation, with 0° denoting tail to head motion. In addition, we presented leftward and rightward motion separately to each eye. To ensure visual stimulation to each eye independently, a thin black bar (0.8 cm) was presented directly underneath the fish. The gratings were locked to the fish orientation in a closed-loop configuration to present the same direction(s) to the fish regardless of its heading orientation. Unless a fish aborted a trial by leaving the active area in the central camera’s field of view, each stimulus was shown for a maximum of 32 seconds. Each experiment ended after 180 minutes. All data was recorded as text files for offline analysis, including frame by frame information about trial structure, x and y position, tail angle, and visual stimulus parameters.

### Simultaneous two-photon calcium imaging and head-fixed tail tracking

To measure neural activity, we performed whole-brain volumetric calcium imaging using a two-photon laser-scanning microscope (Ultima 2P Plus, Bruker, USA). With a pulsed fiber laser (920 nm, Axon, Coherent, USA) as an excitation source, we recorded fluorescent images with a 20X Olympus objective, with an imaging power of 5-10 mW at the specimen. To head-fix larval fish, we immobilized them in 2% weight/volume low melting point agarose (Sigma Aldrich) in embryo medium and removed agarose around the tail for tail tracking. For all experiments, we imaged 712 x 1024 pixels for all functional volumetric records, spanning an area of approximately 470 µm x 660 µm at acquisition rates of 2-4 Hz. For each fish, we recorded ∼20 planes, 7 µm apart, spanning a ∼140 µm volume, lasting 5-6 hours. We started each experiment at the level of the top of the tectum, using the top of Optic tectum and eyes as dorsal landmarks. As in the freely swimming behavior assays, visual stimuli were presented directly underneath the fish. Each stimulus lasted 25 s, beginning with a 20 s stationary period during which the orientation of the grating changed, followed by a motion for 5 s, long enough for neural activity to reach characteristic peaks and return to baseline and allow for behavioral observation associated with each stimulus. All visual stimuli were generated by our custom Python 3.10 software (*pandastim*). Using a 60 Hz, P300 AAXA Pico Projector, only the red LED was connected, allowing for simultaneous visual stimulation and detection of green fluorescence. To track tail movements, we used an infrared camera (Grasshopper3 USB3, Teledyne FLIR LLC, USA), with the tail illuminated by an infrared light (M780L3, ThorLabs, USA) and tracked via *Stytra*^62^ with up to 13 segments at a tracking speed of ∼200 Hz.

### Data processing and analysis

#### Freely-swimming behavior

To process behavioral tracking data, we used our custom Python behavior analysis software (*fishFlux)* for both *ZF* and *DC*. Using the same exclusion criteria for *ZF* or *DC*, fish were removed from further analysis if (1) there were tracking artifacts or (2) the fish did not perform at least one trial of each stimulus. As the smooth swimming patterns in *DC* prevent meaningful bout detection and quantification (**Figure S2**), we calculated behavioral kinematic metrics across the entire time a visual stimulus was presented. To compare *DC* and *ZF* across stimuli, we quantified each fish’s velocity (distance traveled per second), angular velocity (degrees of heading angle per second), and angular acceleration (degrees of heading angle per second squared) and calculated averages or medians across fish. Unless otherwise stated, error bars represent 95% confidence intervals.

#### Two-photon calcium fluorescence source extraction

Following two-photon image acquisition, uncompressed image stacks were corrected for artifact motion using *CaImAn*^35^, an open-source calcium imaging processing library, and its implementation of the NoRMCorre algorithm, which calculates and aligns motion vectors with subpixel resolution. Using *CaImAn* source identification and signal extraction, all further analyses utilized estimated fluorescence traces of signal sources presumed to represent the neural activity of single neurons. As we used transgenic lines with cytosolic expression of *GCaMP*, some sources may represent signals from neural processes. Nonetheless, for simplicity, we refer to these extracted sources as neurons.

#### Visual motion responsive neural activity analysis

Neurons were defined as motion responsive if, for at least one motion direction, their mean activity over 10 s after stimulus onset was above the mean baseline plus 1.8 * standard deviation of the baseline (15 frames directly before stimulus onset) for 80% or more of trials. To compute each neuron’s direction selectivity (DSI), a mean was calculated for responses to the four cardinal (0°, 90°, 180°, 270°) and intercardinal (45°, 135°, 225°, 315°) directions of visual motion. The relative response magnitudes to each stimulus were used to color code response magnitude (intensity) and directionality (hue), resulting in a visual representation of anatomical direction selectivity distribution (see arrow wheel for color code). We used unbiased hierarchal clustering of all motion-responsive neurons using SciPy software^63^ to objectively evaluate functional response types of neurons across the brains of *DC* and *ZF*. Clustering was performed using a responsivity metric (i.e., the difference dF/F between baseline and the stimulus-induced response) for each visual motion cue for all neurons. To quantify the ocular preference of each neuron, we calculated a Binocularity Index (BI) of each neuron:

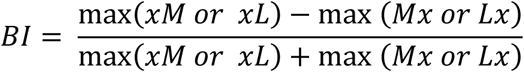

*With xM: right eye medial motion, xL: right eye lateral motion, Mx: left eye medial motion, Lx: left eye lateral motion*

To compare across species and individuals, we calculated a Binocularity index ratio by dividing the relative frequency of binocular neurons (neurons with BI values binned close to 0) by monocular neurons (neurons with BI values binned close to +1 and −1).

#### Tail tracking in head-fixed fish

To correlate neural activity with behavior, we continuously tracked the tail movements of head-embedded fish during calcium imaging using custom software adapted from *Stytra*^62^. As accurate tracking of the translucent, fast-moving fishtail of *DC* required signal-to-noise over time, we only analyzed fish (1) with minimal tracking noise and (2) directed tail behavior in response to visual stimuli. The tail motion was measured by tracking thirteen segments at 200 Hz, yielding a single measured movement variable, ‘tail sum,’ the cumulative angle between neighboring segments. The tail sum standard deviation, calculated over a 100 ms moving window, quantified the increase in tail movement during swims. Each tail event was defined when the tail sum exceeded baseline variance, with an individualized threshold to adjust for variable infrared illumination across fish. To capture *bonafide* tail movements, only tail events ≥100 ms with ≥50 ms intervals between each tail event were included. For each tail event, we calculated the mean tail angle and categorized all events triggered by visual motion stimuli as visually evoked. Principal component analysis (PCA) was conducted on five tail event features: positive and negative mean tail angle, absolute maximum tail angle, duration, and frequency.

#### Motor-associated neural activity analysis

To identify motor decoding neurons across the brain, we developed custom code based on Python SciPy package. We defined bidirectional motor decoding accuracy as the average of the percentage of tail events that are predicted by the dF/F neural activity increase and predict the dF/F neural activity increase. dF/F signal increases were captured by finding peaks across the experiment. We defined a neural activity as motor-associated if the dF/F calcium signal increase coincided with ±0.5 s around each tail event. We also performed motor regressor analysis using Pearson’s correlation. For each fish, neurons with the top 20% of motor decoding accuracy were defined as motor-associated neurons. To understand the behavioral variables contributing to the variations in motor-associated neuron response, the dF/F neural activity of all motor-associated neurons across time was analyzed with principal component analysis. For each tail event, the furthest point of the trajectory from the origin was used to quantify the vigor of the neuronal population. Across all tail events, a Generalized Linear Model (GLM) was used to explain the variation between neuronal population vigor with behavioral variables, including maximal tail angles, tail event durations, tail beat frequency, and number of nearby tail events (±5 s).

To extract turning events in head-fixed fish, we calculated a turn direction index from the mean tail angle across tail events, predicting the neuron response weighted by the magnitude of calcium increase during each event. To pinpoint the spatial location of lateralized turning encoding, the imaging space was divided into 40×40px spatial bins containing corresponding motor-associated neurons. We calculated the average turn direction index for each bin and compared it with a corresponding area on the other side of the midline. The differences in average motor turn direction index across the midline are termed the turning lateralization encoding index. To compare across different individuals, we anatomically aligned maximum intensity projections of each functional imaging plane of each fish to generate the average turning lateralization encoding index in **Figure 4G**.

## Acknowledgments

We would like to thank Drs. Rebecca Yang and Jacob Morra for helpful comments. We thank Drs. Matthew Lovett Barron, Adam Douglass, and the Janelia Fish Facility for gifts of transgenic *DC* and Dr. Misha Ahrens for the *Tg(elavl3:GCaMP7f)* ZF transgenic line. We thank Duke School of Medicine, Jim Burris, Lawerence Frauen for *ZF* and *DC* husbandry, and Sarah Garbatov for assistance during early data collection. Research reported in this publication was supported by the BRAIN initiative of the National Institutes of Health under Award Number RF1NS128895-01. The content is solely the authors’ responsibility and does not necessarily represent the official views of the National Institutes of Health. E.A.N. also was supported by the Whitehall and Alfred P. Sloan Foundation.

## Author contributions

K.E.F. and E.A.N. conceived this project. K.E.F. and M.D.L. designed behavioral and imaging experimental paradigms and analysis pipelines. K.E.F. performed all live *DC* and *ZF* calcium imaging experiments and behavioral experiments. K.E.F. and Z.H. analyzed and compared functional calcium imaging data and behavioral data. K.E.F., Z.H., M.D.L., and E.A.N. wrote the manuscript and contributed to figure preparation.

## Code Availability Statement

All software for behavioral analysis and calcium imaging analysis is available at *github/naumannlab.* All code used for behavior and neural analysis was created using Python and open-source code extensions. The visual stimuli were designed in PANDASTIM available. Due to the enormous size of the imaging data sets, example data will be made available online, and full data sets by request. Instructions for running the code and reproducing published results are available via the code repository README (rendered by GitHub as a web page) and Jupyter Notebooks, which include code to produce specific figures.

## Statistical Analysis

All values are reported as medians unless otherwise stated. Error bars correspond to the distribution. Statistical tests and parameters are specified in the text or figure legends.

## Data Availability Statement

All behavioral data and two-photon calcium imaging data, including processed data as source extracted fluorescence time series, will be publicly available on Duke University’s GitLab server.

## CONTACT FOR REAGENTS AND RESOURCE SHARING

Further information and requests for resources and reagents regarding experimental *ZF* and *DC* data should be directed to Eva Naumann

## Ethics Declaration

None

---

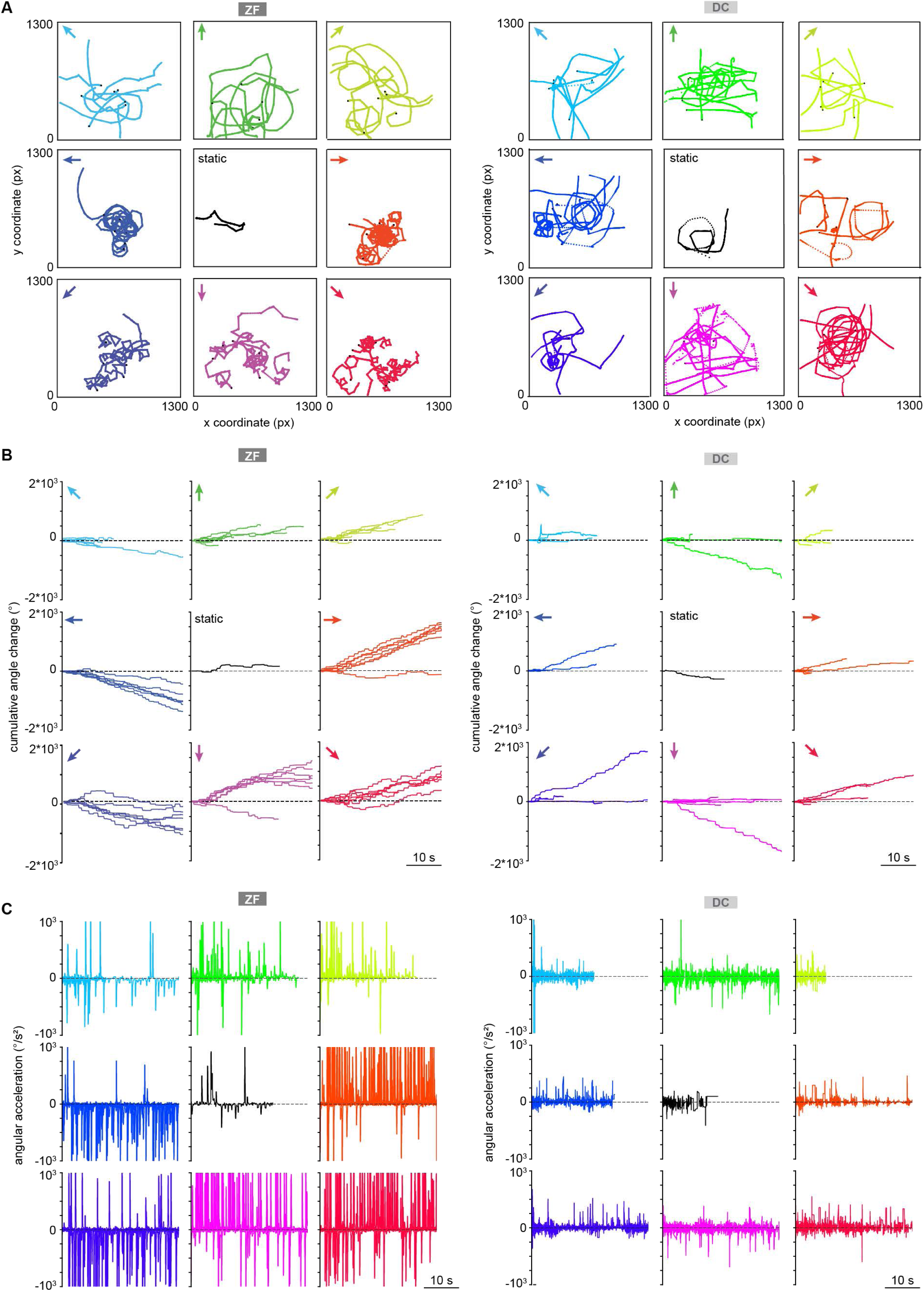

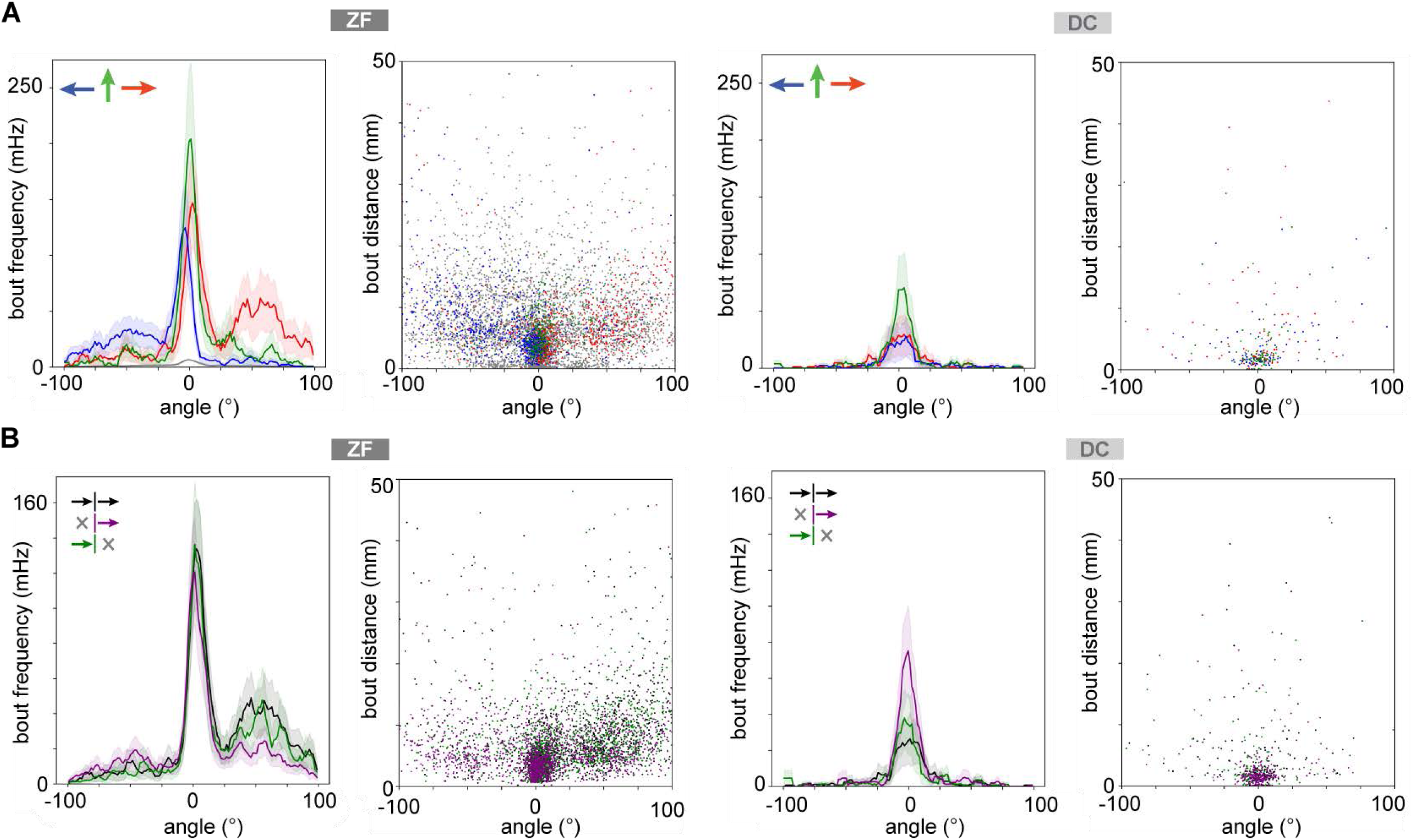

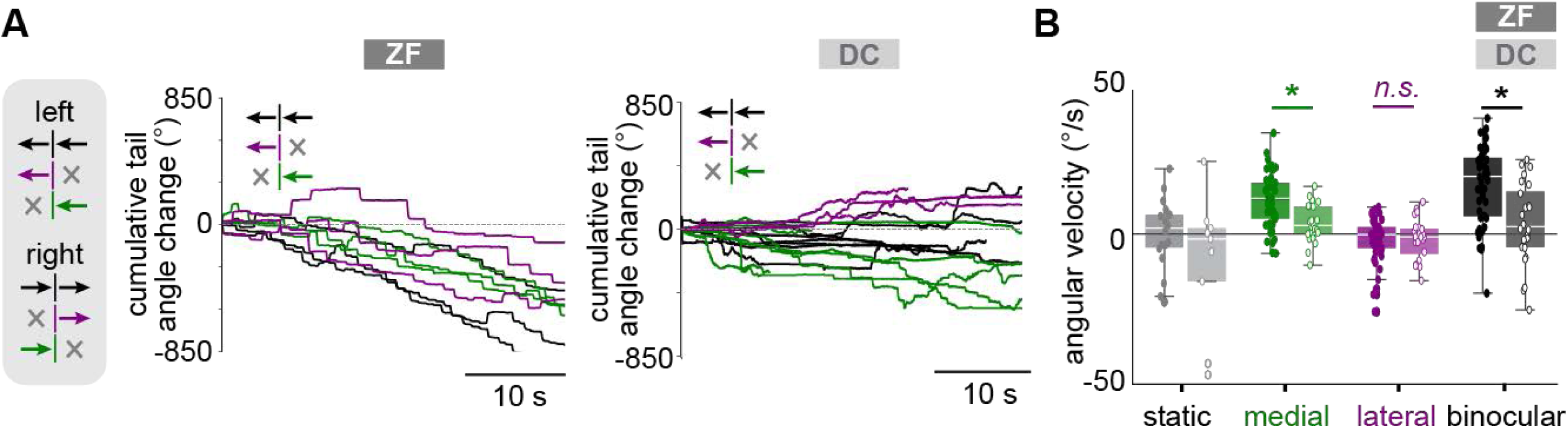

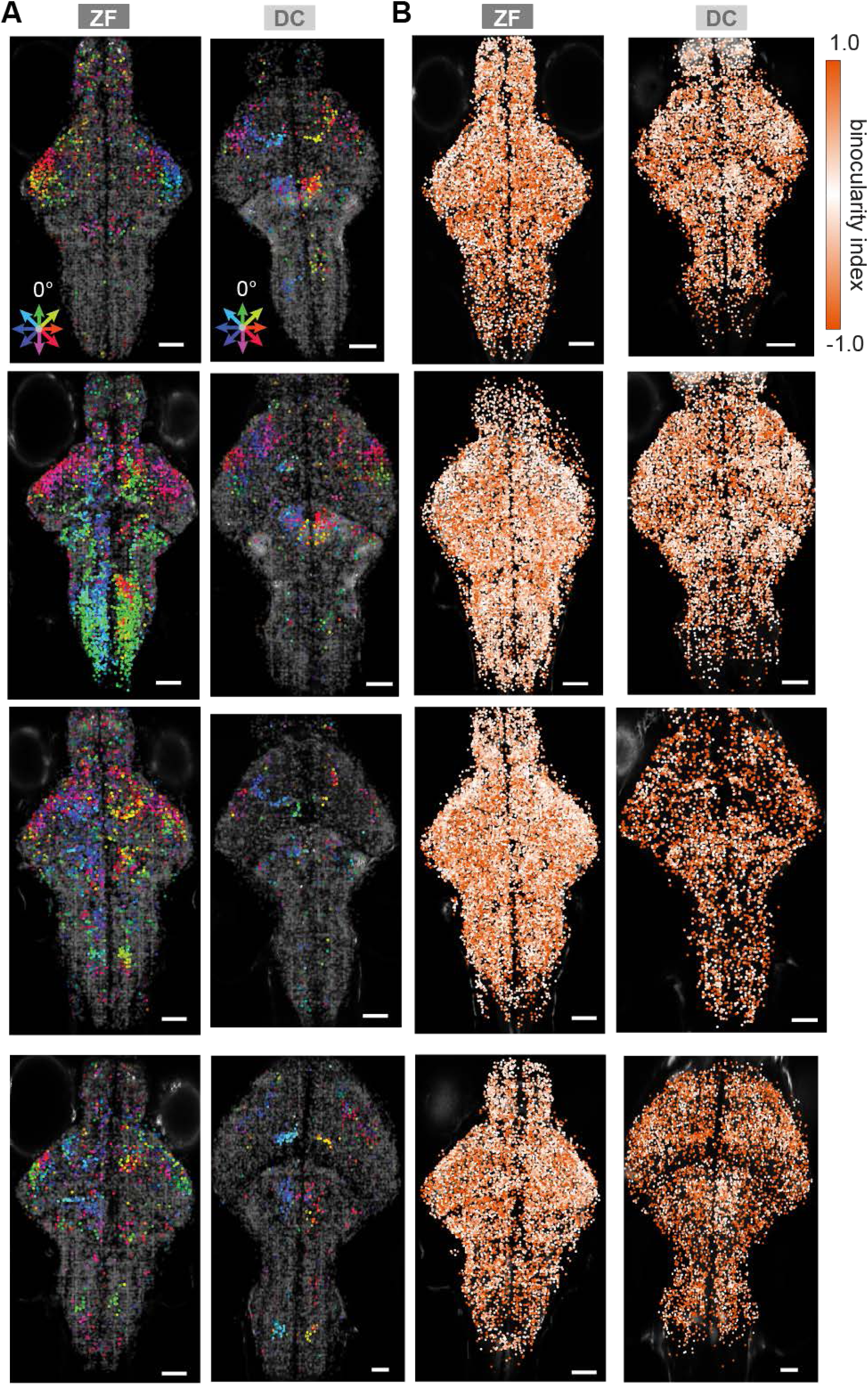

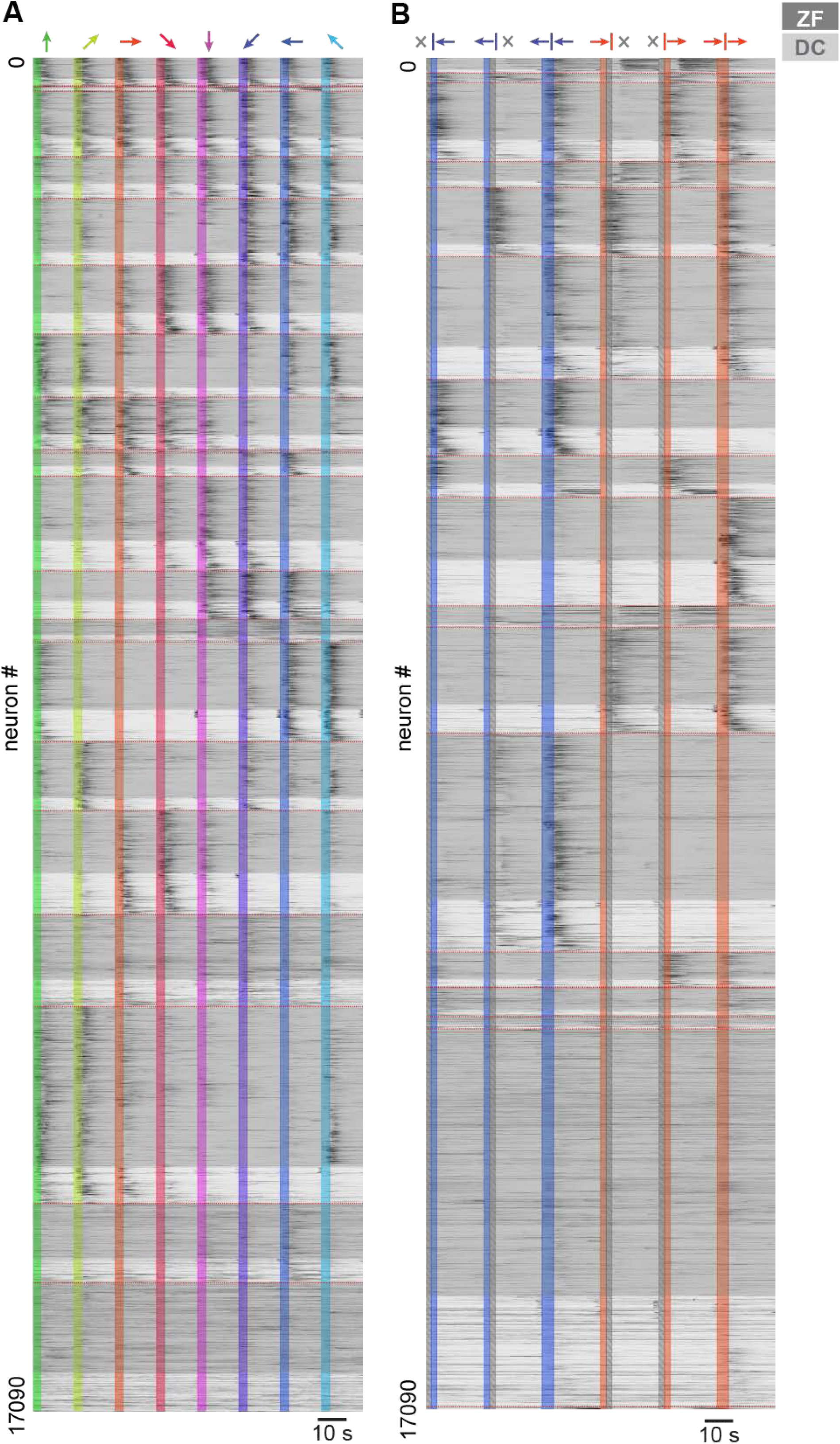

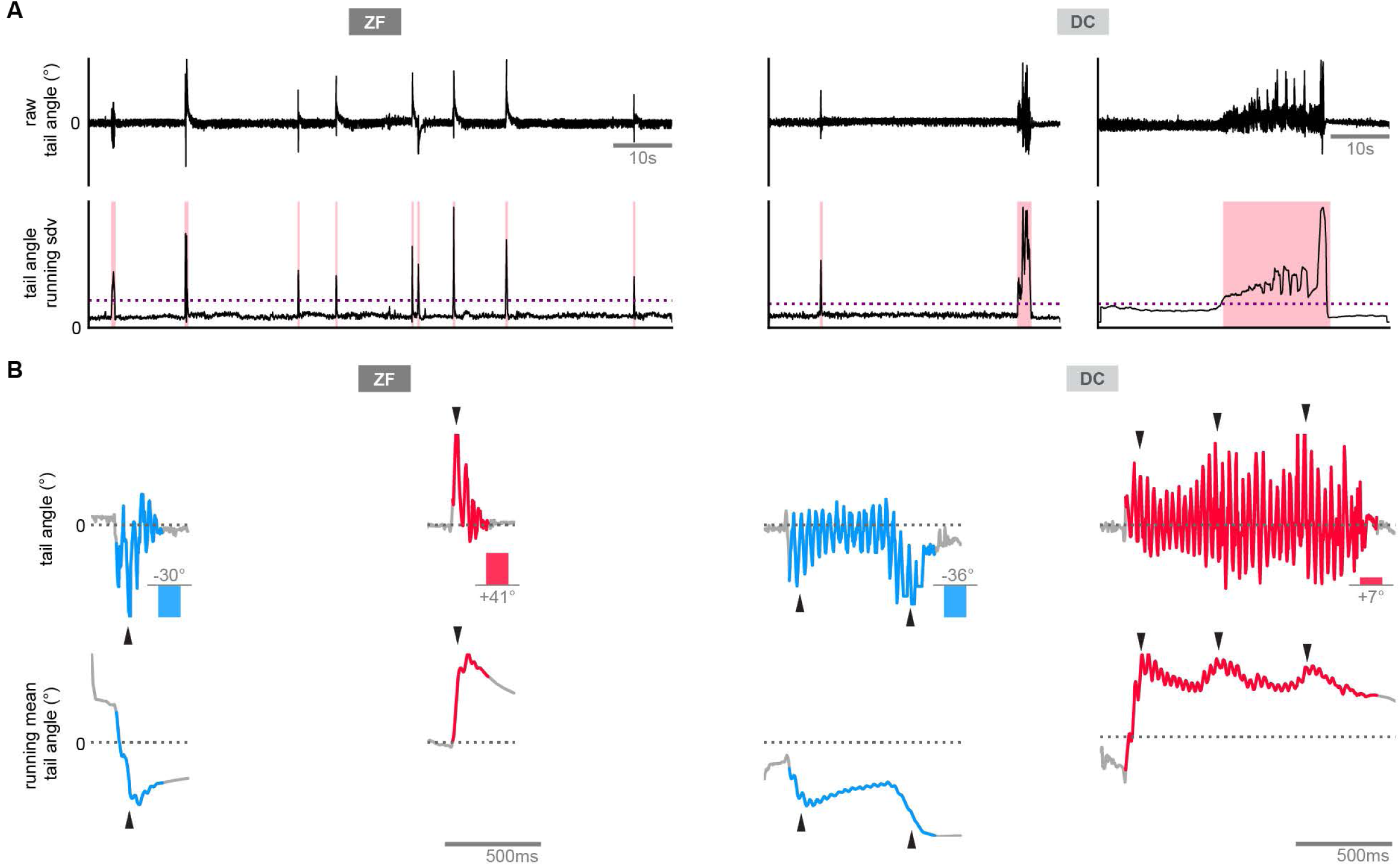

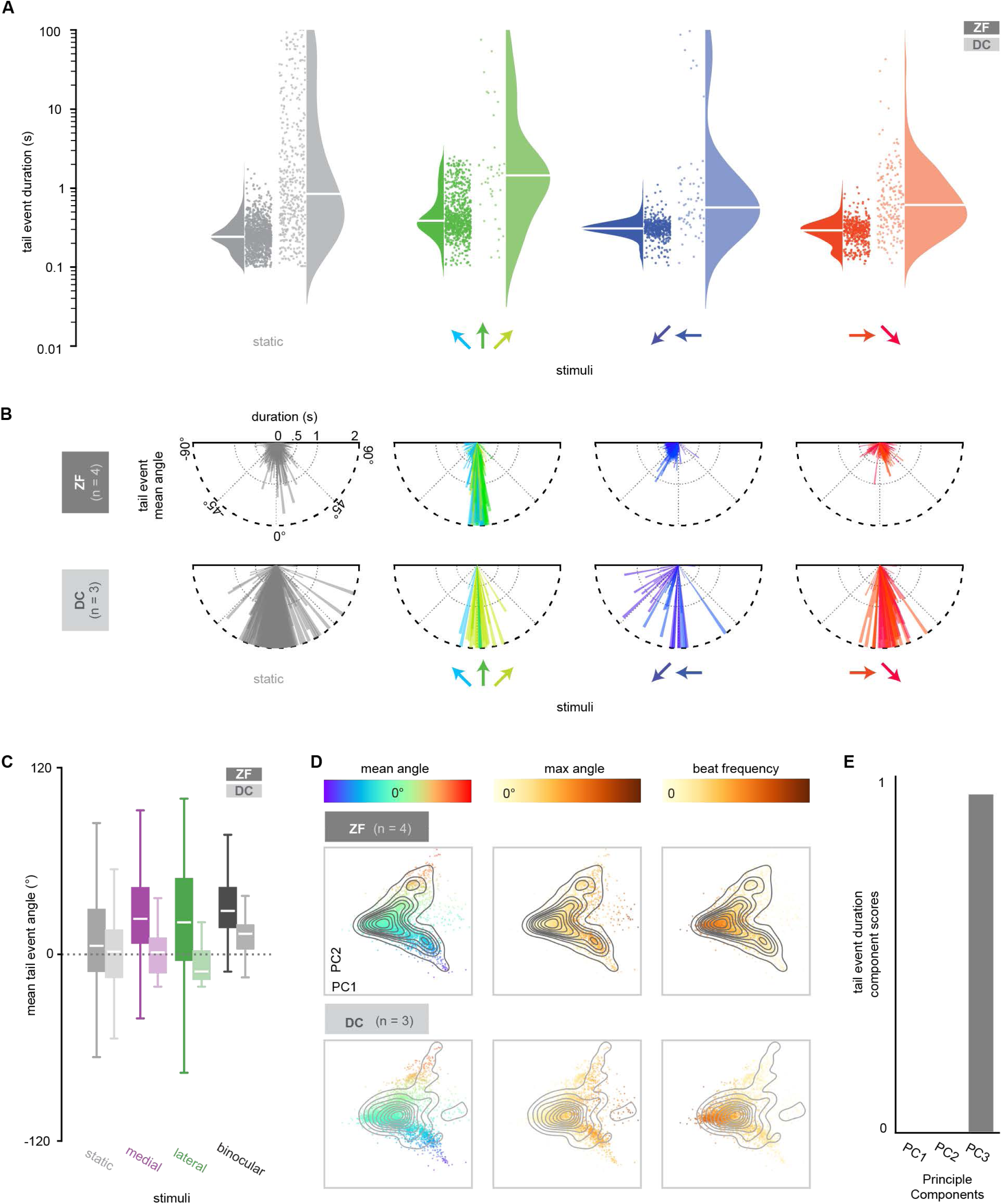

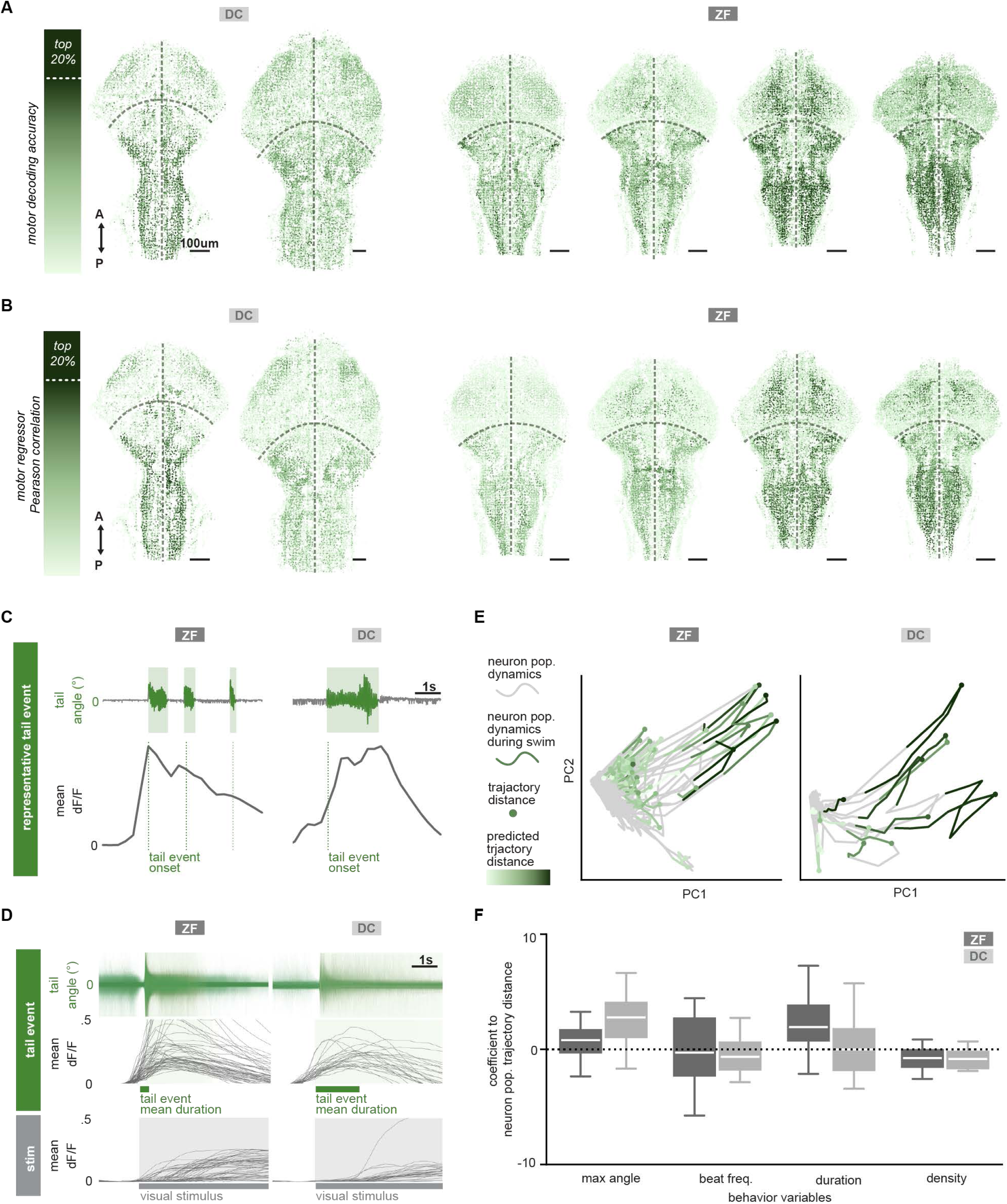

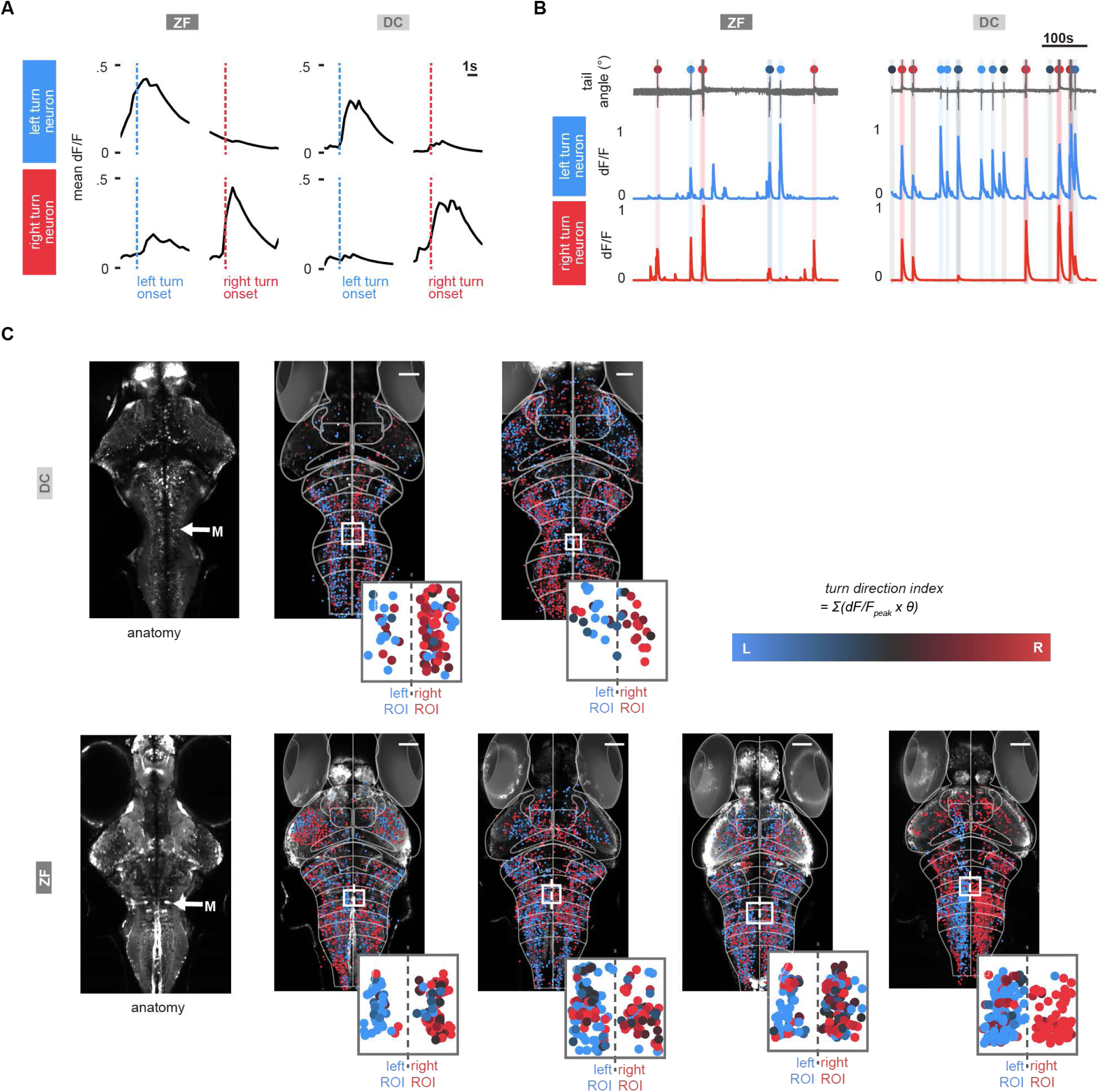

