## Supplementary Information for "Divergent Visuomotor Strategies in Teleosts: Neural Circuit Mechanisms in Zebrafish and *Danionella cerebrum*"

**Figure S1 | OMR behavior of individual larval *ZF* and *DC* (Related to Figure 1).**

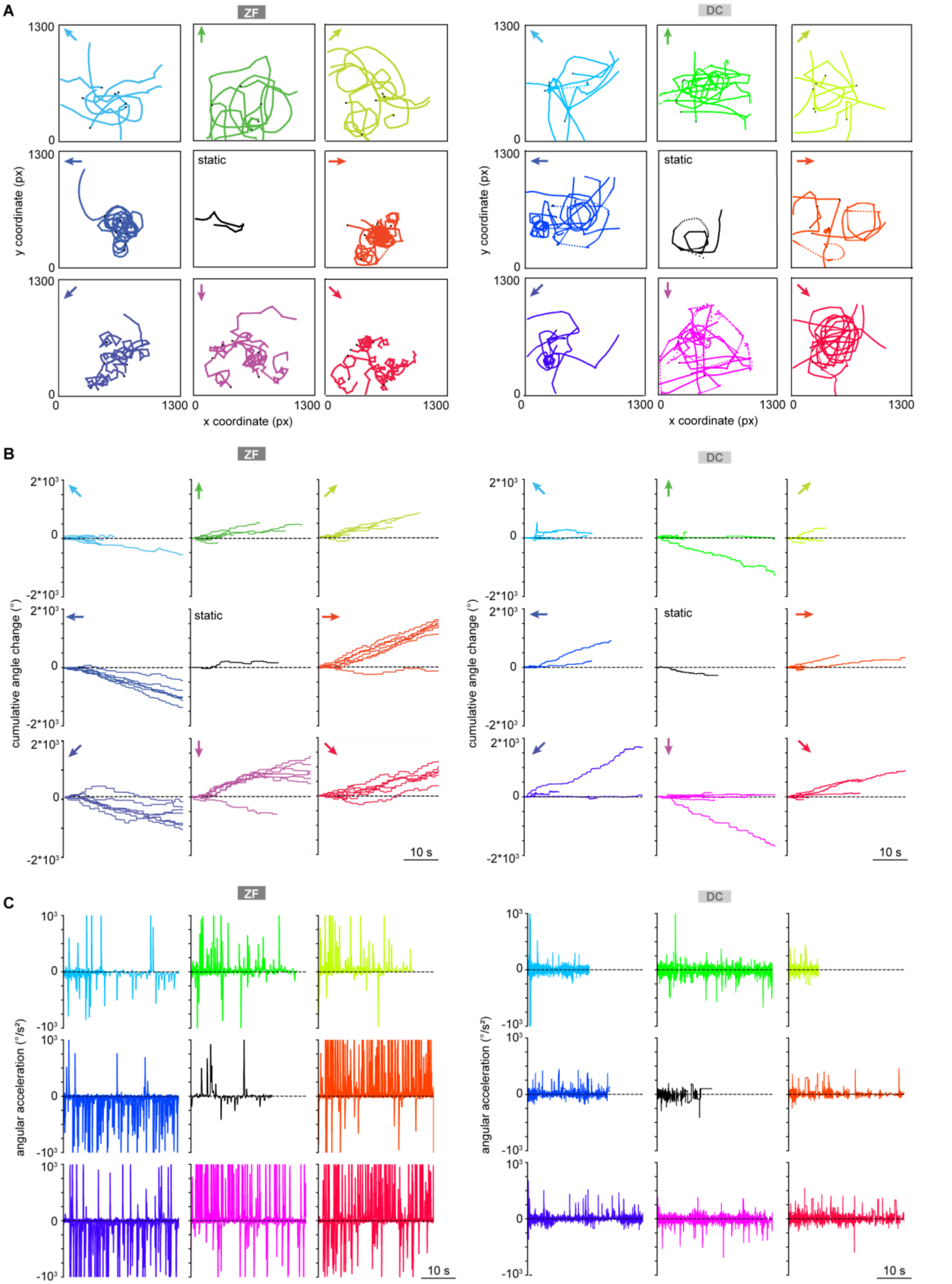

**Figure S1 | OMR behavior of individual larval *ZF* and *DC* (Related to Figure 1).**

**A** Representative x, y position trajectories of *ZF* (left) and *DC* (right) over all trials in response to sinusoidal gratings drifting in eight directions (see color wheel, **Figure 1A**), in addition to a no motion control stimulus (static, black), a stationary 90° sinusoidal grating locked to the fish's body axis. Each line represents one trial of motion-induced behavior; each line is colored according to the stimulus direction presented during that trial (see colored arrows). The starting positions for each trial are marked with black dots. Notice that *DC* trials are shorter in duration due to the more frequent premature abortion of OMR trials, as individual *DC* tended to exit the 'active' field of view. *ZF* display tight spiraling turning behavior when presented with non-forward motion patterns, while *DC* shows less stimulus modulation.

**B** Cumulative angle change over time in response to the same trials as in **A**. The 'staircase' shape of many traces indicates that both *DC* and *ZF* exhibit abrupt changes in direction, indicating that both species use punctuated orientation adjustments.

**C** Angular acceleration of behavior during the same trials as in **A**, **B**.

**Figure S2 | *ZF* and *DC* bout-wise kinematic analysis (Related to Figure 1).**

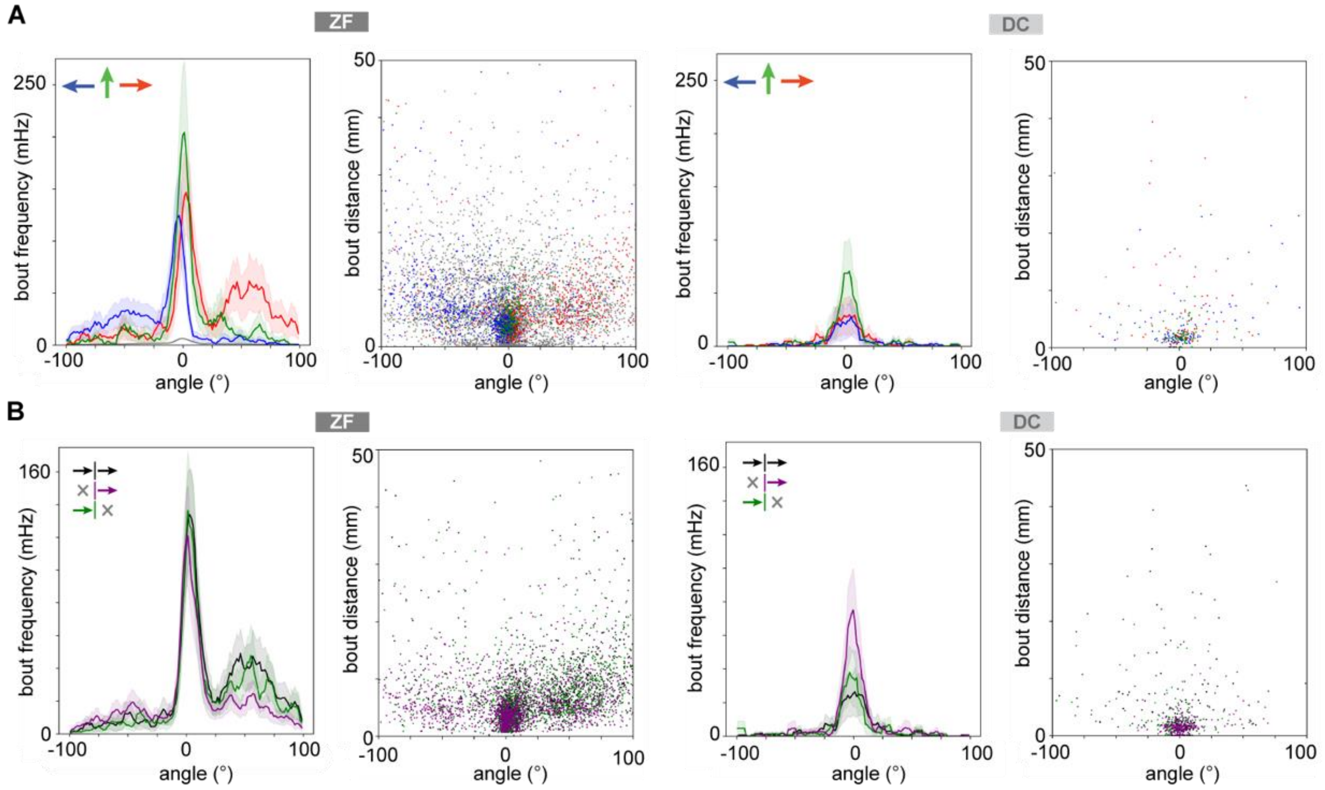


**Figure S2 | *ZF* and *DC* bout-wise kinematic analysis (Related to Figure 1).**

**A** Average histograms of absolute frequency and distance per bout angle change across *ZF* and *DC* (N = *ZF*: 12 fish, *DC*: 12 fish) over all trials in response to forward (green), leftward (blue), and rightward (red) moving gratings, and no motion control stimulus (static, grey). Bouts are identified by velocity peaks using *Stytra*[62]. *ZF* perform frequent and directional bouts following motion cues, as previously reported[10], compared to *DC.* Shaded error is SEM. Applying the same bout-wise kinematic analysis in *DC* yields fewer bouts with lower heading direction changes. Due to *DC* smooth continuous swimming patterns, kinematic bout analysis is inadequate to describe *DC* OMR behavior. Nonetheless, *DC* are capable of following optic flow (**Video 1**).

**B** Same analysis as in **A**, for behavior trials during monocular medial, lateral and binocular left and rightward motion, collapsed for direction. *ZF* show directional bouts, as previously reported[10], while *DC* show few non-directional bouts.

**Figure S3 | Monocular driven OMR behavior in *ZF* and *DC*. (Related to Figure 1).**

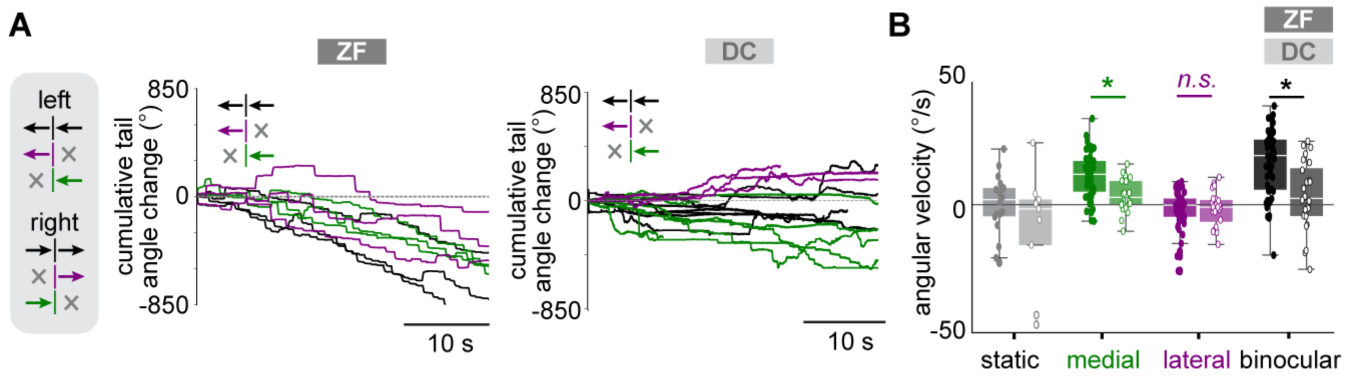


**Figure S3 | Monocular driven OMR behavior in *ZF* and *DC* (Related to Figure 1).**
**A** *Left*, schematic of monocular and binocular motion stimuli that were presented to *ZF* and *DC*. Arrows indicate the direction of eye-specific motion cues, i.e., medial motion towards the body midline, lateral motion away from the body midline, and binocular motion shown in the same direction to both eyes. *Right*, Cumulative heading direction angle changes during monocular medial (green), monocular lateral (purple), and whole field left motion (black) for representative *ZF* and *DC*.

**B** Median angular velocity across *DC* and *ZF* during the presentation of static, monocular, and binocular moving stimuli collapsed for each direction. Positive angular velocity indicates 'correct' turns, following the stimulus direction. Both species respond most strongly to binocular stimuli, but *DC* consistently exhibits slower angular velocity than *ZF* (p> 0.05, two-way ANOVA).

**Figure S4 | Whole-brain visual motion correlation and binocularity maps (Related to Figure 2).**

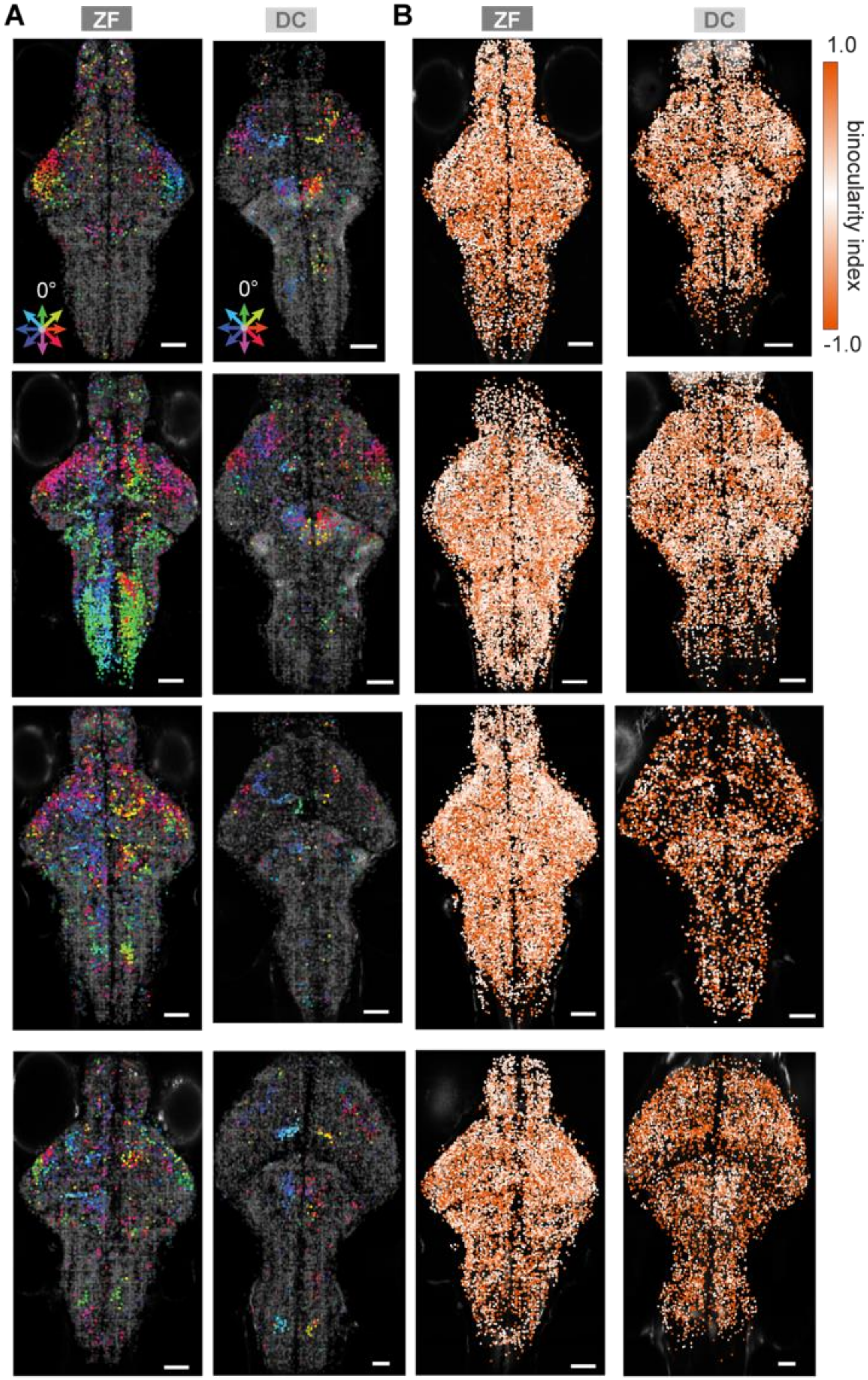

**Figure S4 | Whole-brain visual motion correlation and binocularity maps (Related to Figure 2).**

**A** Matched individual *ZF* and *DC* whole-brain motion correlation direction-responsive activity maps (*cf*., **Figure 2D**). Each dot represents a neural source, with its hue indicating direction preference (see colored arrow wheel), plotted on a greyscale *GCaMP* image. Only strongly direction-selective neurons with weighted mean response >0.2 dF/F are colored by direction selectivity index (DSI), otherwise grey. Scale bar, 100 um.

**B** Binocularity maps for the same fish in **A**. Each neuron is color-coded based on its binocularity index (BI), with neurons responding exclusively to the left or right eye colored orange (BI > +/-0.6). Neurons respond equally to left and right eye (BI = 0).

**Figure S5 | Shared functional types of motion-responsive neurons in *DC* and *ZF* (Related to Figure 2).**

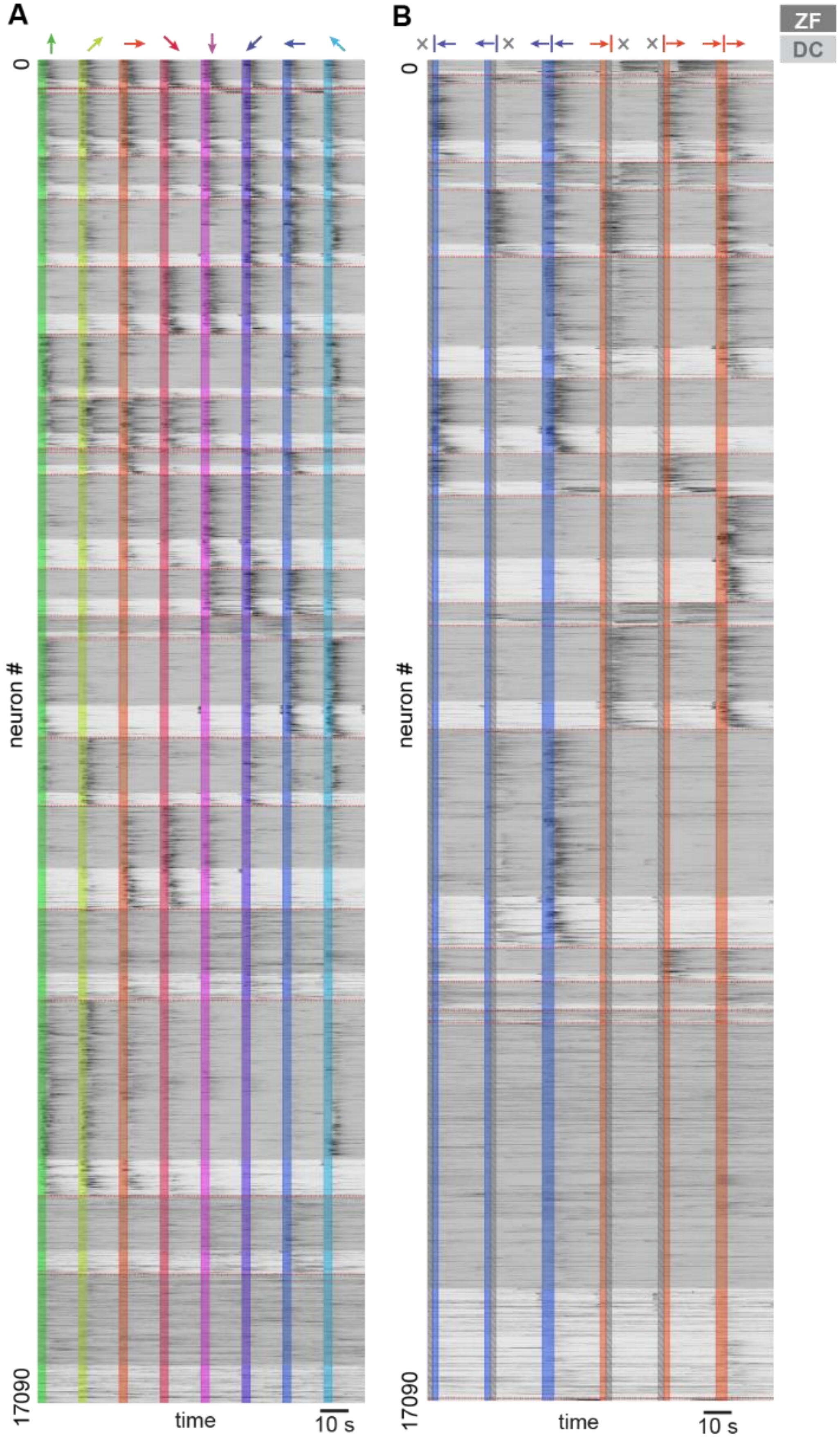

**<u>Figure S5 | Shared functional types of motion-responsive neurons in *DC* and *ZF* (Related to Figure 2).</u>**
**A** Heatmap of average dF/F responses for all motion-responsive neurons with a weighted mean dF/F above 0.2 across stimulus repetitions from all recordings of *ZF* and *DC*. Hierarchical clustering functionally classified neurons based on each neuron's responses to eight whole-field motion stimuli, revealing that all clusters contain neurons from both species (*ZF* neurons shaded in dark grey, *DC* in light grey). Notably, *ZF* show a higher prevalence of forward-tuned neurons, mirroring *ZF* strong increased behavioral drive in response to forward motion in comparison to *DC*. Stimulus direction is indicated by arrows at the top, with colored bars marking stimulus onset. **B** Average dF/F heatmaps of neurons from **A**, re-clustered based on their responses to monocular and binocular stimuli (indicated by symbols at the top). Most neurons across species exhibit lower activity in response to monocularly presented stimuli, but all clusters contained neurons from both species.

**Figure S6 | Tail movement tracking analysis in head fixed in *DC* and *ZF* (Related to Figure 3).**

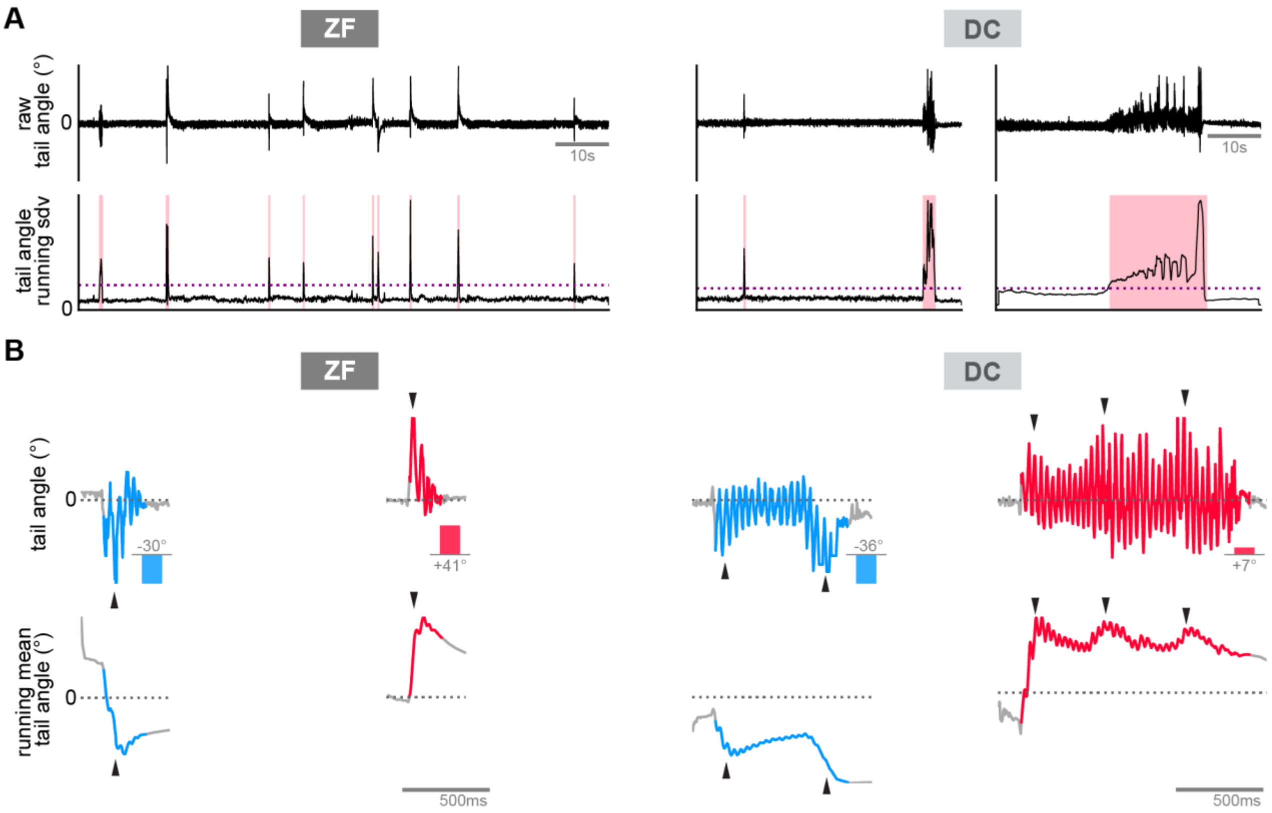


**Figure S6 | Tail movement tracking analysis in head fixed in *DC* and *ZF* (Related to Figure 3)**

**A** Snapshot of tail event extraction for *ZF* (left) and *DC* (right). The upper trace shows the raw tail angle sum across tracking frames, while the lower trace displays tail angle variation as the standard deviation in a moving window across time. Red-shaded areas show detected tail events, isolated using a standard deviation threshold (purple dashed line) above baseline tracking variance. *ZF* exhibit short, burst-like tail events, whereas *DC* show many long periods of tail movement that can be detected by our algorithm, even head-fixed *DC.*

**B** Example left and right turn tail event in *ZF* (left) and *DC* (right). *Top*, representative, extracted left (blue) and right (red) turn events, with inset bar graph showing heading direction change. *DC* performs turns with sharp angle changes (black arrowheads) interspersed within straight swimming events (tail angles ~ 0°). In contrast, *ZF* typically perform a single, distinct sharp turn at the beginning of the bout. Bottom, running mean tail angle across time.

**Figure S7 | Analysis of head-embedded behavior *ZF* and *DC* in response to OMR stimuli (Related to Figure 3).**

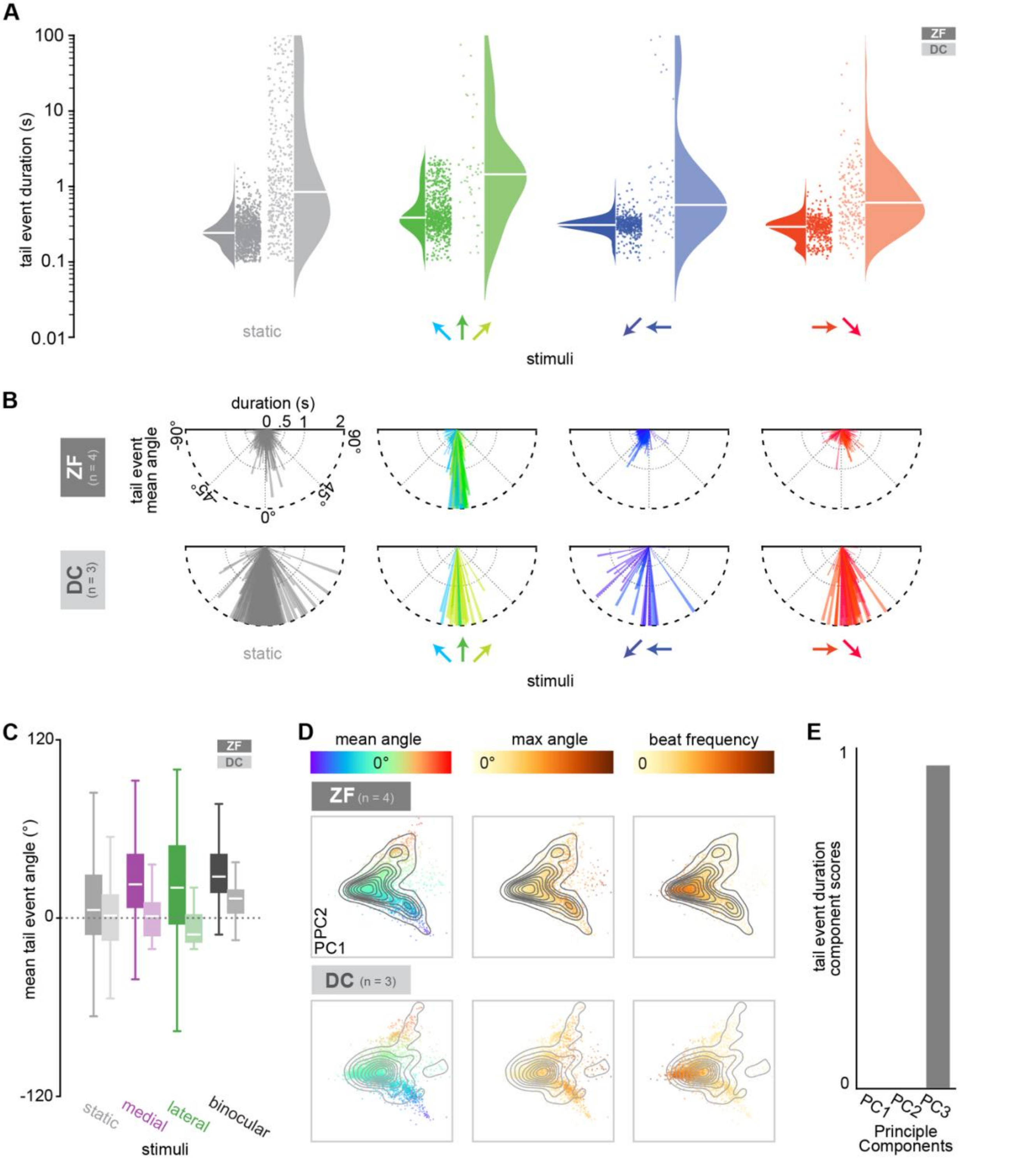

**<u>Figure S7 | Analysis of head-embedded behavior *ZF* and *DC* in response to OMR stimuli (Related to Figure 3).</u>**

**A** Distribution of all tail event durations for *ZF* and *DC* during visual stimulation. White lines indicate medians. Each dot indicates a tail event. For *ZF*, tail events during forward, forward left, and forward right stimuli last significantly longer than those during other directional stimuli (Two-way ANOVA, Tukey post hoc, * $p < 0.05$). In *DC*, tail events are less lateralized and do not show significant shortening for oblique, turn-inducing stimuli.

**B** Distribution of all tail events for head-embedded *ZF* (N = 4 fish, 3568 events) and *DC* (N = 3 fish, 855 events). Each bar represents a tail event, with bar direction reflecting tail angle and color indicating visual motion stimulus presented during the tail event (see color wheel). Bar length represents the duration of the tail event, capped at 2 seconds. As in freely swimming *ZF* (**Figure 1D**), *ZF* bout durations are significantly modulated, with forward, forward left, and forward right stimuli driving significantly longer bouts than oblique, directional turn inducing stimuli (Two-way ANOVA, Tukey post hoc, * $p < 0.05$). *DC* tail events show no significant shortening for oblique stimuli.

**C** Box plots of the mean tail angle across all monocular and binocular left and right visual stimuli for *ZF* and *DC*, white lines represent the medians. *DC* displayed far fewer directed bouts during monocular stimulation trials in comparison to *ZF*, mirroring behavior observed in freely swimming *DC*.

**D** PCA analysis of tail event captured in head-embedded *ZF* (N = 4 fish, 3568 events) and *DC* (N = 3 fish, 855 events). Each dot represents one tail event, color-coded by mean angle (left), absolute max angle (middle), and frequency (right). Despite differences in tail event durations (**E**), *DC* and *ZF* show similar overall distribution. However, *DC* exhibit fewer sharp turns and lower frequency with larger tail angle change during straight swim events due to *DC* performing interspersed sharp turns during straight swims.

**E** In the PCA analysis (**D**), tail duration variations are captured by PC3, which were not depicted in (**D**).

**Figure S8 | Motor-associated population encoding of behavioral variables in *ZF* and *DC* (Related to Figure 4).**

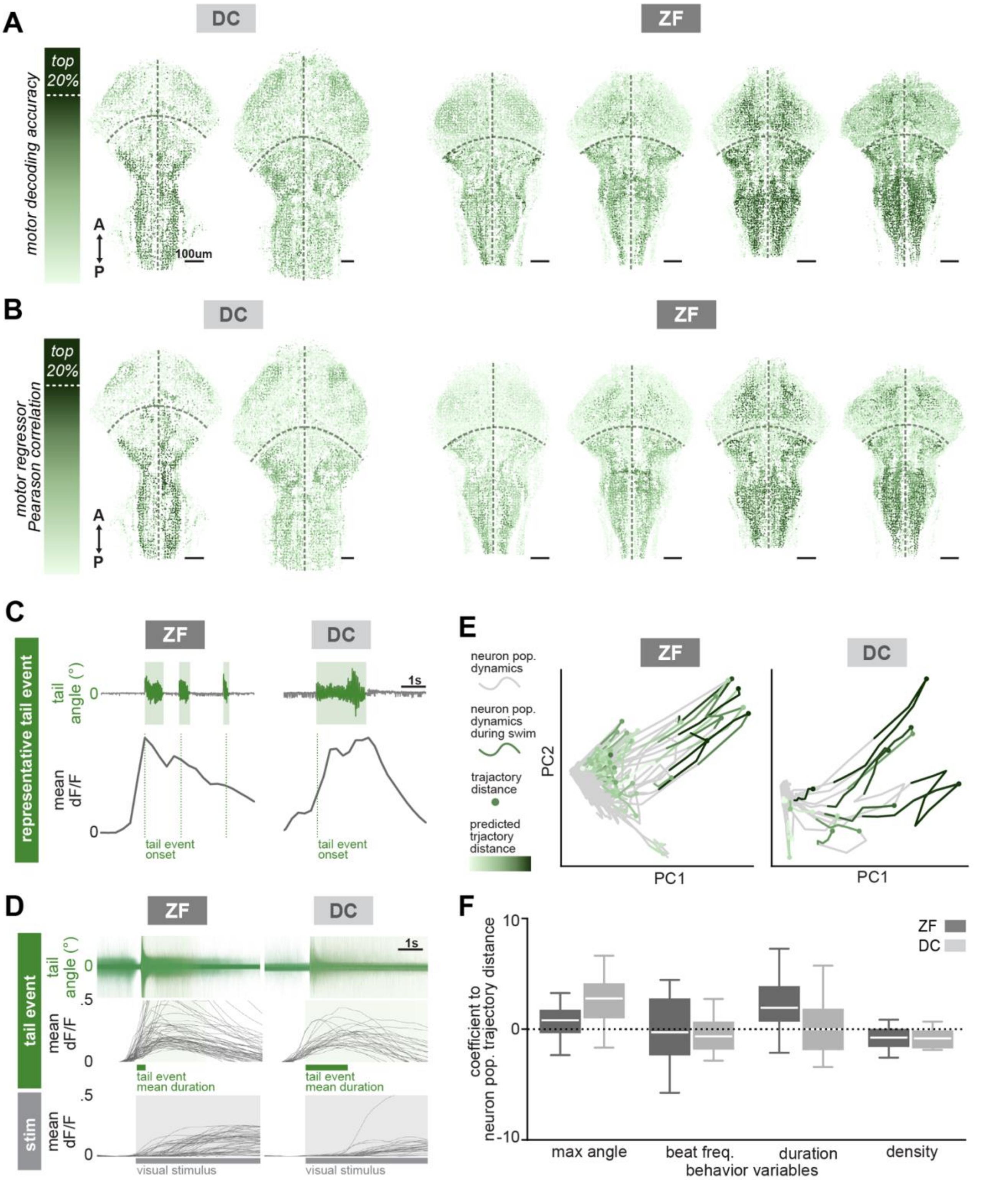

**Figure S8 | Motor-associated population encoding of behavioral variables in *ZF* and *DC* (Related to Figure 4).**

**A** Motor decoding accuracy across all neurons in two additional *DC* and *ZF* brain volumes. Top motor-associated neurons are found in both species in the hindbrain. Darker colors indicate higher motor decoding accuracy.

**B** Similar motor-associated neurons are extracted based on Pearson's correlation coefficient, with the motor regressor representing all swim events. Motor decoding accuracy in (**A)** extracts highly similar neurons from this motor regressor analysis. Darker colors indicate neurons with higher Pearson's correlation coefficient.

**C** Representative tail events showing the neuron dynamics during temporally coupled bouts in *ZF* and sustained swim event in *DC*. *Top*, representative tail angle. *Bottom*, average calcium activity from the entire population of motor-associated neurons, with dashed lines marking the start of each tail event.

**D** Activity of top 20% motor-associated neuron population aligned around swim onset (N = ZF: 4 fish, 14063 neurons; DC: 2 fish, 5790 neurons). *Top*, swim events aligned to onset, *middle*, average calcium activity across all motor-associated neurons aligned to the start of all swim events, bottom, aligned to visual motion stimulus onset.

**E** Dynamics of representative motor-associated neuron population during imaging, with swim event periods (from swim onset to swim end +2 s) shaded in green. A Generalized Linear Model (GLM) was applied to identify behavior variables driving neuron population dynamics. Dot positions represent the furthest PCA trajectory distance from the origin, used as the proxy for swim-driven population dynamics. Darker shades of green color indicate stronger neuron population dynamics predicted from the GLM model.

**F** GLM results demonstrate the contributions of different behavior variables to motor-associated neuron dynamics. Each neuron population resides in one imaging plane because tail traces are only associated with one two-photon imaging z-plane. In *DC*, maximal angle primarily drives motor associated-neuron dynamics, while in *ZF*, swim duration plays a greater role, contributing substantially to the differences in motor associated-neuron dynamics.

**Figure S9 | Medial hindbrain neurons correlate with turning behavior in both species (Related to Figure 4).**

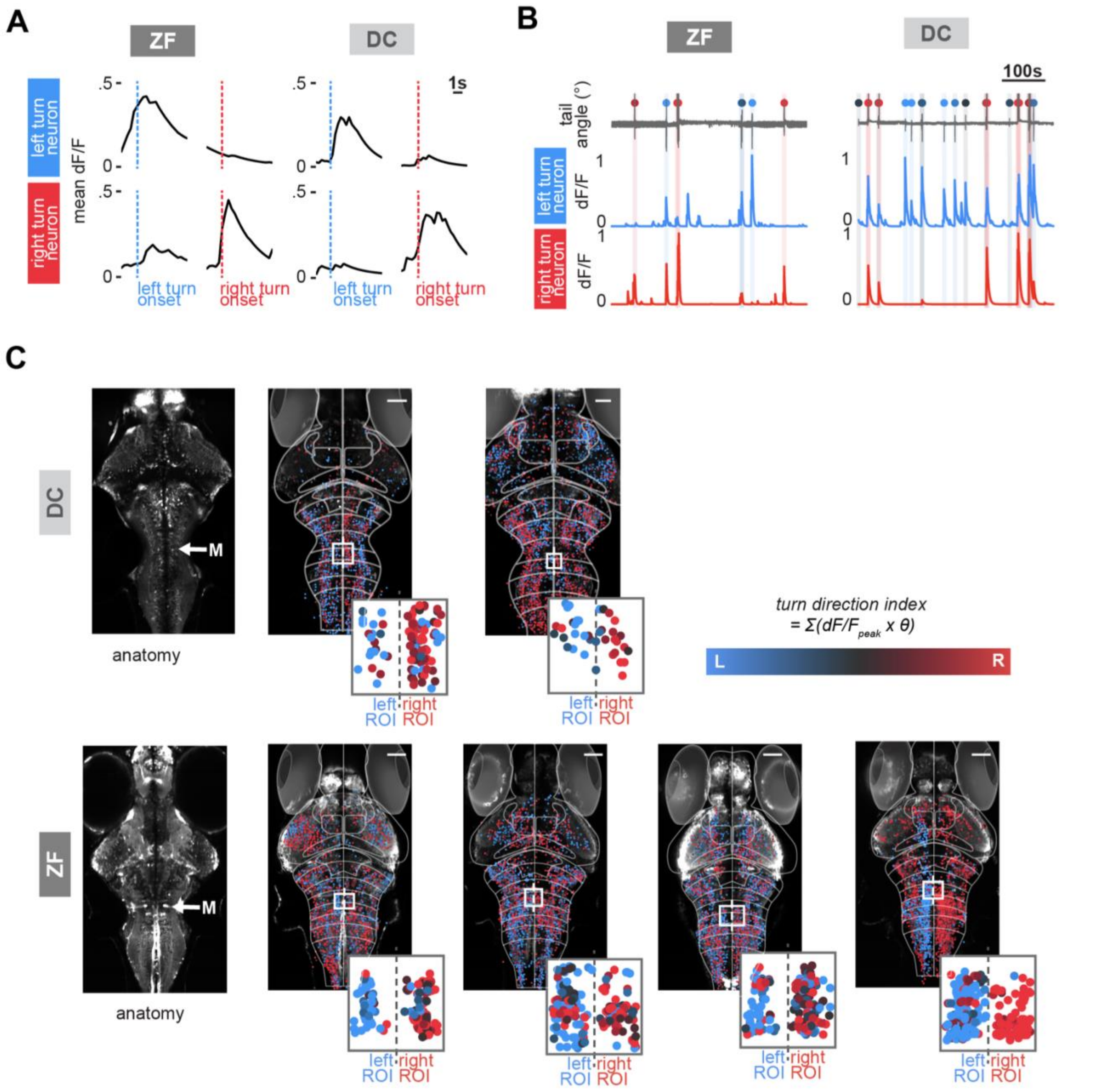


**Figure S9 | Medial hindbrain neurons correlate with turning behavior in both species (Related to Figure 4).**

**A** *Top*, average dF/F neural activity of a representative *ZF* and *DC* neuron associated with leftward turning during left and right turns. Dotted lines mark the onset of the left (blue) and right (red) turn bout. *Bottom*, average dF/F calcium for representative right turning- associated neurons in *ZF* and *DC.* The strong turn direction associated neural activity of these neurons suggests that they may be participating in controlling left or rightwards turns.

**B** *Top*, raw tail angle, with detected left (blue dots) or right (red dots) turns for *ZF* and *DC*. Below dF/F traces show neural activity during these bouts. Representative rightward and leftward turn-associated neurons were chosen from left and right ROI depicted in **C.** Blue and red shaded vertical areas indicate the onset and duration of right and left bouts.

**C** *Left*, maximum intensity projection image from a two-photon plane in a representative *ZF* and *DC* with *GCaMP7f* expression. White arrow points to the location of Mauthner cells in the hindbrain, between rhombomeres 4 and 5, which were used to anatomically identify the squared ROIs to the right. *Right*, top 20% motor-associated neurons in all *ZF* and *DC* color-coded by the turn direction index calculated for each individual neuron. Zoom-ins show the enlargement of boxed ROIs from the top plot.

**Video Legends:**

**<u>Video 1 | *DC* performs OMR by swimming continuously, interspersed with sharp angle turns.</u>**

This video shows representative behavioral trials of a larval *Danionella cerebrum (DC)* performing OMR while presenting forward, right, left, and backward moving gratings, capturing 5-7 seconds. The left display shows the visual stimulus presented to the fish, and the right display shows the tracked fish overlaid with a red dot to indicate the fish's head position and a blue line marking the tracked tail. Notice the continuous tail beating and sharp angle turns when *DC* responds to leftward, rightward, backward moving gratings. Related to **Figure 1**.

**<u>Video 2 | Anatomical flythrough of a 7-day-old *Tg(neurod1:GCaMP7f) Danionella cerebrum*.</u>**

Video of *GCaMP7f* expression across the brain of a representative *Tg(neurod1:GCaMP7f) DC* panning through 350 μm volume (50 planes, 7 μm apart) dorsal to ventral. Related to **Figure 2**.

**<u>Video 3 | Whole-brain activity recording and tail tracking during visual stimulation.</u>**

This video shows examples of raw calcium imaging and behavioral tracking in head-embedded, tail-free *DC* and *ZF* while presenting visual stimuli from below. Each video clip shows the ongoing moving grating visual stimulus (left), raw two-photon fluorescence for a single plane across the brain (middle), and infrared highspeed video recording (right), with the blue line indicating the central body axis, the red line shows the tracked tail segments used to calculate the tail sum angle (below). Related to **Figure 2-4**.
